# OmniCellAgent: An AI Scientist for Omic-Driven Scientific Discovery

**DOI:** 10.1101/2025.07.31.667797

**Authors:** Di Huang, Hao Li, Wenyu Li, Heming Zhang, Tianqi Xu, Yifei Lu, Kaiwen Fang, Zixi Xu, Justin Chen, Patricia Dickson, Marco Sardiello, William Buchser, Jonathan Cooper, Carlos Cruchaga, Pirooz Eghtesady, Guangfu Li, S. Peter Goedegebuure, David DeNardo, Li Ding, Ryan C. Fields, Ming Zhan, J Philip Miller, Michael Province, Yixin Chen, Philip Payne, Fuhai Li

## Abstract

Real-world biomedical scientific discovery operates as an iterative lifecycle integrating four core pillars: the targeted identification and analysis of question-specific omics datasets; the context-aware interpretation of molecular data using rich biomedical prior knowledge database; comprehensive literature reviews; and the subjective creativity and expert intuition of human scientists. Together, these pillars drive robust evidence synthesis and novel hypothesis generation. The recently reported AI agents support automated omics analysis and literature review, which typically require users to predefine and curate disease-specific datasets, which is a process that remains challenging and time-consuming. In this study, we present OmniCellAgent, a novel multi-agent AI framework built on large-scale single-cell RNA sequencing (scRNA-seq) datasets, and can autonomously retrieve and analyze disease and control-related scRNAseq datasets of diverse cell types across tissues and conditions. Moreover, it incorporates a biomedical prior knowledge and literature review agents, and disease domain-specific expert agents to systematically annotate omic data-derived targets. By aggregating evidence across agents, the framework generates structured analytical reports and potential scientific hypotheses. We evaluated OmniCellAgent across multiple disease settings, demonstrating its ability to identify relevant datasets, generate omic data analysis results, and produce structured reports and scientific hypotheses. Our findings demonstrate that multi-agent AI systems lower the technical barriers to omics-driven research, thereby accelerating scientific discovery and hypothesis generation in biomedical research and precision medicine. The source code is publicly available at: https://github.com/FuhaiLiAiLab/OmniCellAgent.

---

Omics-driven research has become central to this process, enabling systematic characterization of disease-specific molecular targets and signaling pathways that can guide personalized therapies. Advances in next-generation sequencing have produced unprecedented growth in large-scale omics datasets across tissues, diseases and experimental conditions. For example, The Cancer Genome Atlas (TCGA) [50, 55] was the first project to systematically generate multi-omics profiles for tens of thousands of cancer samples across diverse cancer types [12]. The Cancer Cell Line Encyclopedia (CCLE) project [13] generated multi-omics data from 1,000 cancer cell lines. In the cancer Dependency Map project (DepMap) [1, 51] and the Connectivity Map project (CMAP) [22, 23, 46], genome-wide genetic and large-scale compound perturbations were performed in these cancer cell lines to investigate genetic targets and drugs that regulate cancer cell growth. Following these cancer-focused projects, omics datasets have also been generated for other diseases, including Alzheimer’s disease (AD) [4, 25], across many studies [16, 25, 32, 53] and resources such as the Religious Orders Study and Memory and Aging Project (ROSMAP) [3, 26, 42]. In the Long-Life Family Study (LLFS) project [57], omics datasets have been used to identify protective biomarkers and signaling pathways that promote healthy aging and exceptional longevity. More recently, single-cell omics technologies have enabled high-resolution characterization of intra- and inter-cellular signaling networks across heterogeneous cellular microenvironments. For example, single-cell omics data for AD have been generated by projects such as the Brain Cell Atlas [6] and are available through the AD Knowledge Portal [10, 19] or the CZ CellxGene repository [37]. The Human Tumor Atlas Network (HTAN) [8, 41] generates single-cell omics data from diverse cancer samples to study tumor microenvironment signaling evolution. The Human Cell Atlas (HCA) [40] includes scRNA-seq data from more than 70 million cells across different diseases. Moreover, the Arc virtual cell atlas recently released Tahoe-100M, a dataset of 100 million single-cell transcriptomic profiles designed to study transcriptomic variation caused by 1,100 small molecules across 50 cancer cell lines [63]. Finally, the Biohub Billion-Cell Project aims to generate billion-scale single-cell datasets, and its first dataset of approximately 22 million primary human CD4+ T cells perturbed by genome-scale CRISPR was recently released [64]. These rapidly expanding single-cell omics resources are valuable for decoding complex cell-specific signaling systems in disease microenvironment (ME).

However, omics-driven scientific discovery remains a fundamental challenge in biomedical research. Discovery is an iterative process: scientists identify, integrate, annotate and analyze relevant omics datasets; interpret results using prior knowledge from biomedical databases and comprehensive literature review; and then generate testable hypotheses about molecular mechanisms, therapeutic targets and drug combinations. This process is subjective, labor-intensive and difficult to scale [14, 20, 27, 30, 43, 59]. To support biomedical discovery, recent work has explored agentic AI systems that combine reasoning, tool use and domain-specific knowledge. These systems coordinate specialized agents for tasks such as data analysis, literature retrieval and hypothesis generation in autonomous or semi-autonomous workflows. The year 2025 has been marked by the rise of AI agents: autonomous systems capable of perceiving, reasoning and acting within an environment for scientific discovery [54, 59]. For example, Gottweis et al. [14] introduced AI co-scientist, a multi-agent system that proposes and refines research hypotheses through an autonomous loop over large-scale literature reports. Biomni [20] was developed as a general-purpose AI agent for biomedical data analysis. Kosmos [30] analyzes user-uploaded omics datasets and performs literature search. OmniScientist [43] analyzes large-scale literature cohorts and generates novel hypotheses. Similarly, scientists have envisioned biomedical AI agents that integrate with experimental platforms and use prior knowledge to drive discovery while keeping humans in the loop [11, 31]. Dynamic tool use [38], where agents flexibly call external knowledge and text-mining utilities, enables systems to handle complex queries spanning the full spectrum of biomedical data. A recent review systematically examined newly reported agentic AI models and autonomous AI agents designed for scientific discovery [54, 59]. Despite these advances, several key challenges remain for omics-centered multi-agent AI frameworks. First, existing systems often assume that users have already selected, retrieved, and formatted disease-specific omics datasets; this requirement presents a major bottleneck for many researchers and severely limits cross-disease data reuse. Second, current workflows rely heavily on isolated literature reviews or pre-uploaded data, failing to dynamically integrate raw single-cell omics data analyses, structured biomedical knowledge graphs, targeted literature evidence, and broader web context into a single reproducible scientific report. Third, most platforms generate narrative hypotheses or descriptive summaries rather than the full spectrum of artifacts required for rigorous inspection and validation, such as processed omics tables, retrieved evidence logs, synthesized graphs, and standalone interactive visualizations. Fourth, existing agents often lack the capacity to seamlessly annotate omics-derived targets with prior biomedical knowledge and literature, a critical step for elucidating dysfunctional biological processes and signaling pathways.

In this study, we present OmniCellAgent, a multi-agent AI framework designed to address these challenges by integrating large-scale single-cell RNA sequencing (scRNA-seq) data, biomedical prior knowledge and literature-driven reasoning. Built on a comprehensive collection of scRNA-seq datasets spanning diverse tissues, diseases and cell types, OmniCellAgent autonomously retrieves and analyzes relevant disease-control datasets in response to user queries. The framework integrates tools that operate on complementary forms of evidence: raw and processed OmniCellTOSG omics data handled by bioinformatic pipelines [62]; biomedical knowledge-graph evidence derived from BioMedGraphica and related graph resources [58, 60, 61]; peer-reviewed literature retrieved through PubMed; and broader, timely web evidence retrieved through Google search. Its outputs are also structured multimodal reports to facilitate AI human collaboration for scientific discovery: the system returns structured reports, reusable analysis tables, retrieved and summarized evidence, graph-based annotations and standalone interactive visualizations. By unifying data retrieval, statistical analysis, biomedical graph reasoning, literature search, web search and expert synthesis within a single agentic framework, OmniCellAgent extends the “omni” scope of previous biomedical agents to cell-type-specific omics discovery. We demonstrate its utility across multiple disease contexts, highlighting its ability to reduce barriers to data access, enhance omic-data analysis interpretability and accelerate hypothesis generation in precision medicine. The framework is released as an open-source resource to allow users to build upon and further improve it to better support the broader biomedical research community.

## RESULTS

### Overall architecture and workflow of OmniCellAgent

OmniCellAgent uses LangGraph [24] to coordinate a planner-executor-reporter workflow (Figure 1). The planner converts a user query into an ordered research plan, the executor invokes specialized agents for omics analysis, knowledge-graph retrieval, literature search, web search and expert synthesis, and the reporter integrates their outputs into an evidence-grounded report. A persistent `AgentState` records the plan, intermediate results, structured data fields and process log, allowing downstream agents to reuse compact evidence rather than reparse long conversational traces. Figure 2 illustrates an example state flow for an Alzheimer’s disease query and shows the example payloads produced at each stage.

**Fig. 1.**
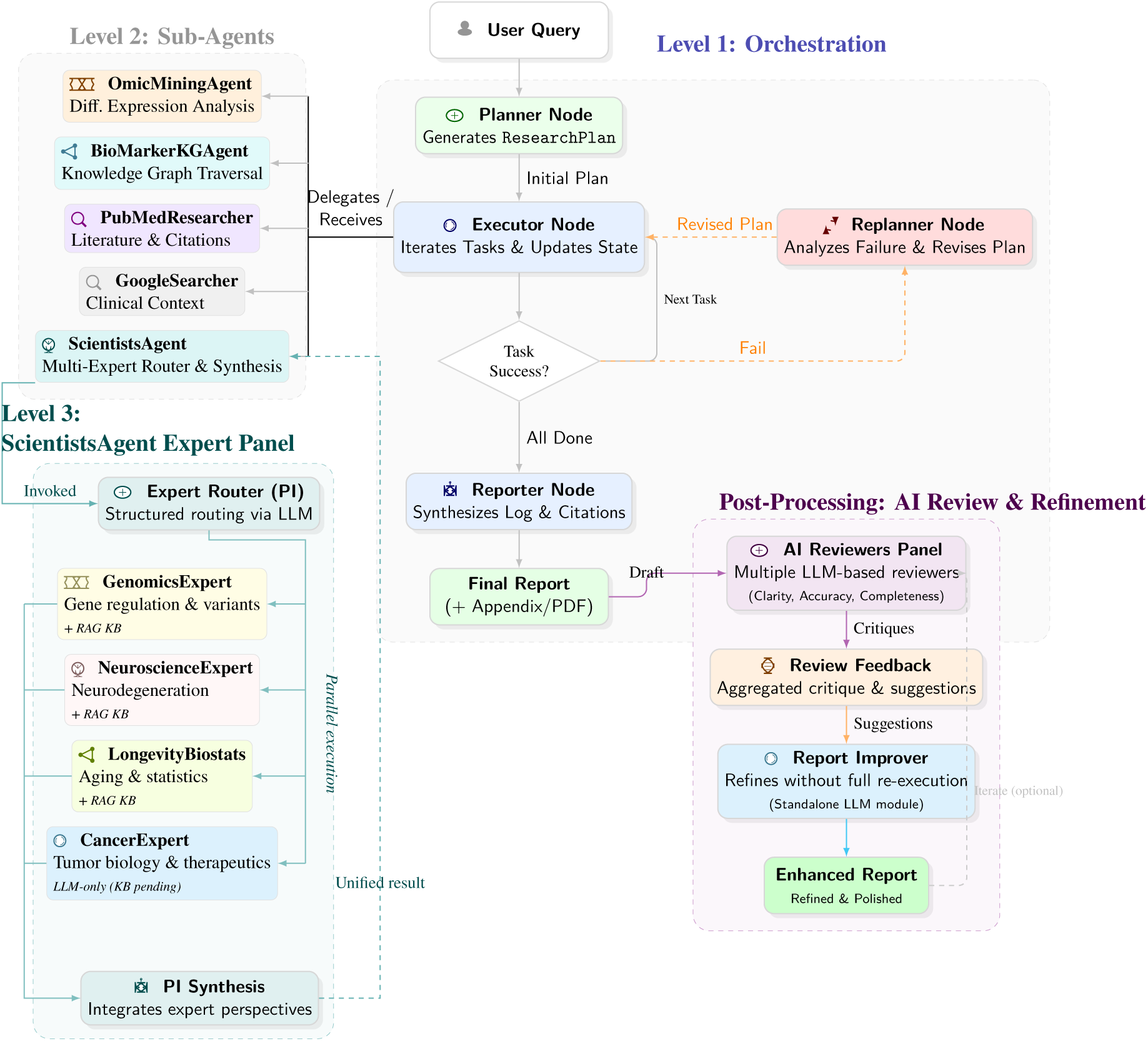
Three-level architecture of OmniCellAgent with post-processing refinement. **Level 1 (Orchestration):** A Planner decomposes the user query into a `ResearchPlan`; the Executor dispatches each task to the appropriate sub-agent, a Decision node triggers the Replanner on failure, and the Reporter compiles the final synthesis with citations. **Level 2 (Sub-Agents):** Five specialised sub-agents—OmicMiningAgent, BioMarkerKGAgent, PubMedResearcher, GoogleSearcher, and ScientistsAgent—are invoked in parallel or sequentially by the Executor. **Level 3 (Expert Panel):** Inside ScientistsAgent, an Expert Router (PI) performs structured LLM-based routing to four domain experts (Genomics, Neuroscience, Longevity/Biostatistics, Cancer). **Post-Processing (AI Review & Refinement):** After the initial report is generated, an AI Reviewers Panel evaluates the draft for clarity, accuracy, and completeness. The aggregated feedback is passed to a standalone Report Improver module that refines the report without re-executing the full agent system, producing an enhanced final output with optional iterative refinement.

**Fig. 2.**
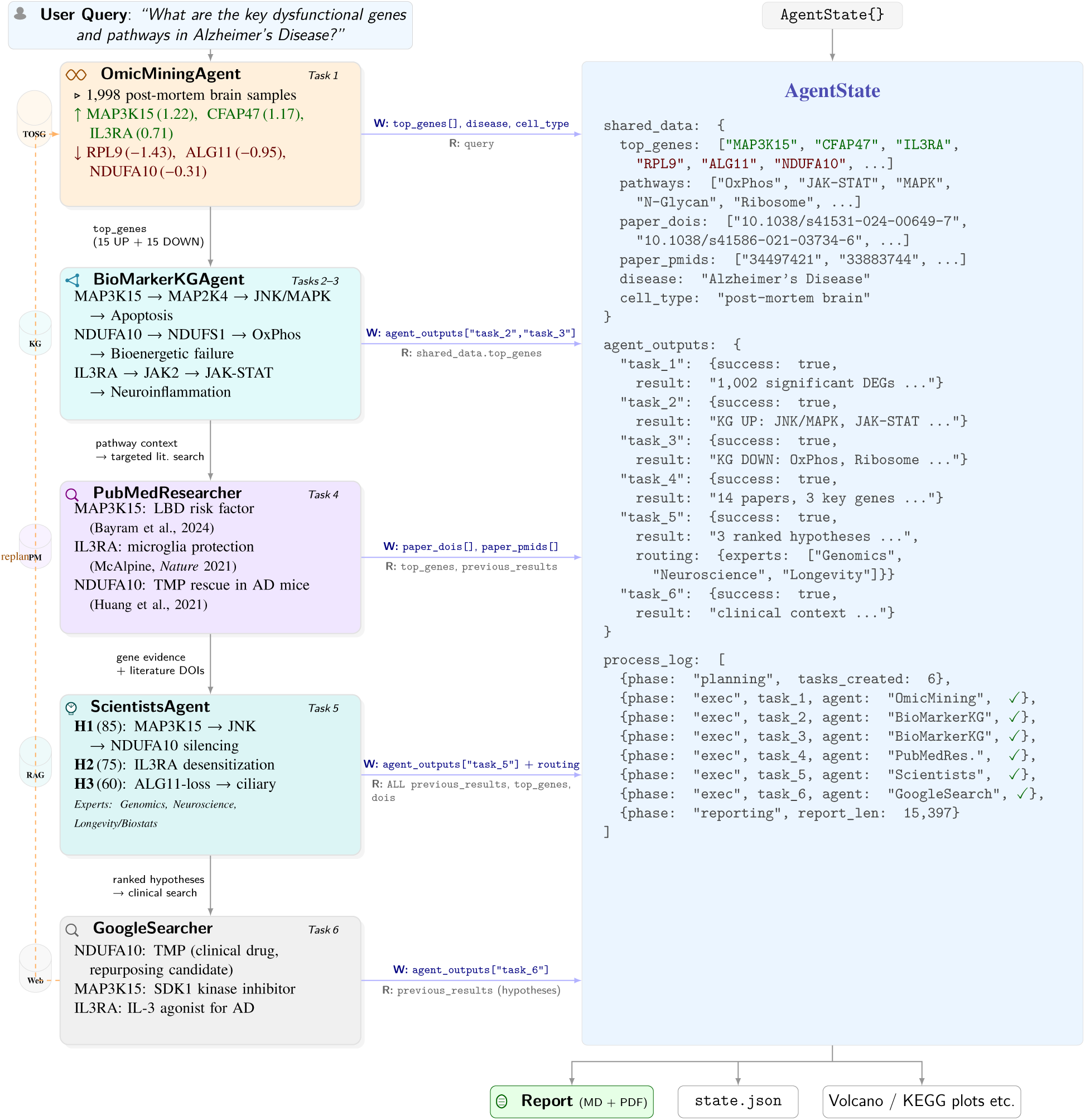
Example agent workflow for an Alzheimer’s Disease query using the OmniCellAgent system. The Planner decomposes the query into six sequential tasks; each sub-agent receives the accumulated `AgentState` and writes its outputs back. **Read/write annotations** on each horizontal arrow show the specific state fields consumed and produced: OmicMiningAgent populates `top_genes` (1,002 DEGs from 1,998 post-mortem brain samples), BioMarkerKGAgent maps those genes to pathways via PrimeKG, PubMedResearcher adds literature DOIs for the top hits, ScientistsAgent routes to three domain experts (Genomics, Neuroscience, Longevity/Biostats) and synthesizes ranked mechanistic hypotheses, and GoogleSearcher assesses translational viability (e.g. TMP for NDUFA10 rescue, SDK1 for MAP3K15 inhibition). The downward arrows indicate how each step leverages results from the preceding step—enabling progressively deeper reasoning that mirrors the workflow of a human research team. The following sections have close-ups of the system to illustrate the contribution of each agent in this example realization.

This state-centered design separates rich conversational context from compact, programmatically accessible evidence. Omics analysis writes ranked genes and pathways, literature search writes DOI/PMID identifiers, and expert synthesis writes ranked hypotheses and validation ideas. The same state object supports replanning after failed subtasks and preserves provenance for the final report and appendix, keeping long multi-agent runs modular and traceable.

### Case studies of using OmniCellAgent

We evaluated OmniCellAgent across three different disease settings to assess its utility for practical, multi-disease, scRNA-seq-data-driven case studies—namely Alzheimer’s disease (AD), pancreatic ductal adenocarcinoma (PDAC), and lung adenocarcinoma (LUAD). Specifically, in AD, we performed two complementary comparisons, diseased versus healthy and male versus female within the AD cohort, while in PDAC and LUAD we focused on disease-versus-control contrasts. Across all analyses, we applied a consistent workflow including differential expression testing, correlation structure analysis of top genes, dimensionality reduction and group separation assessment using PCA and PERMANOVA, and pathway enrichment analysis. Together, these results show that the agent can recover robust transcriptomic differences at both the gene and pathway levels, capture biologically meaningful population-level separation between conditions, and generalize across multiple disease contexts, with the AD case further demonstrating its ability to reveal both disease-associated and sex-associated molecular heterogeneity.

### Transcriptomic profiling identifies convergent inflammatory and bioenergetic signatures in Alzheimer’s disease

Differential expression analysis of a large post-mortem brain transcriptomic cohort comparing Alzheimer’s disease (AD) and control samples identified broad molecular changes consistent with neuroinflammation, impaired mitochondrial function, and altered protein homeostasis. In the analyzed dataset, the most prominent upregulated genes included MAP3K15, CFAP47, several long non-coding RNAs, and IL3RA, whereas the most prominent downregulated genes included NDUFA10, ALG11, and RPL9. The report summarizes a total sample size of 1,998 and highlights widespread transcriptomic disruption in AD relative to controls (Figure 3; sex-stratified AD analysis is provided in the Supplementary Information).

**Fig. 3.**
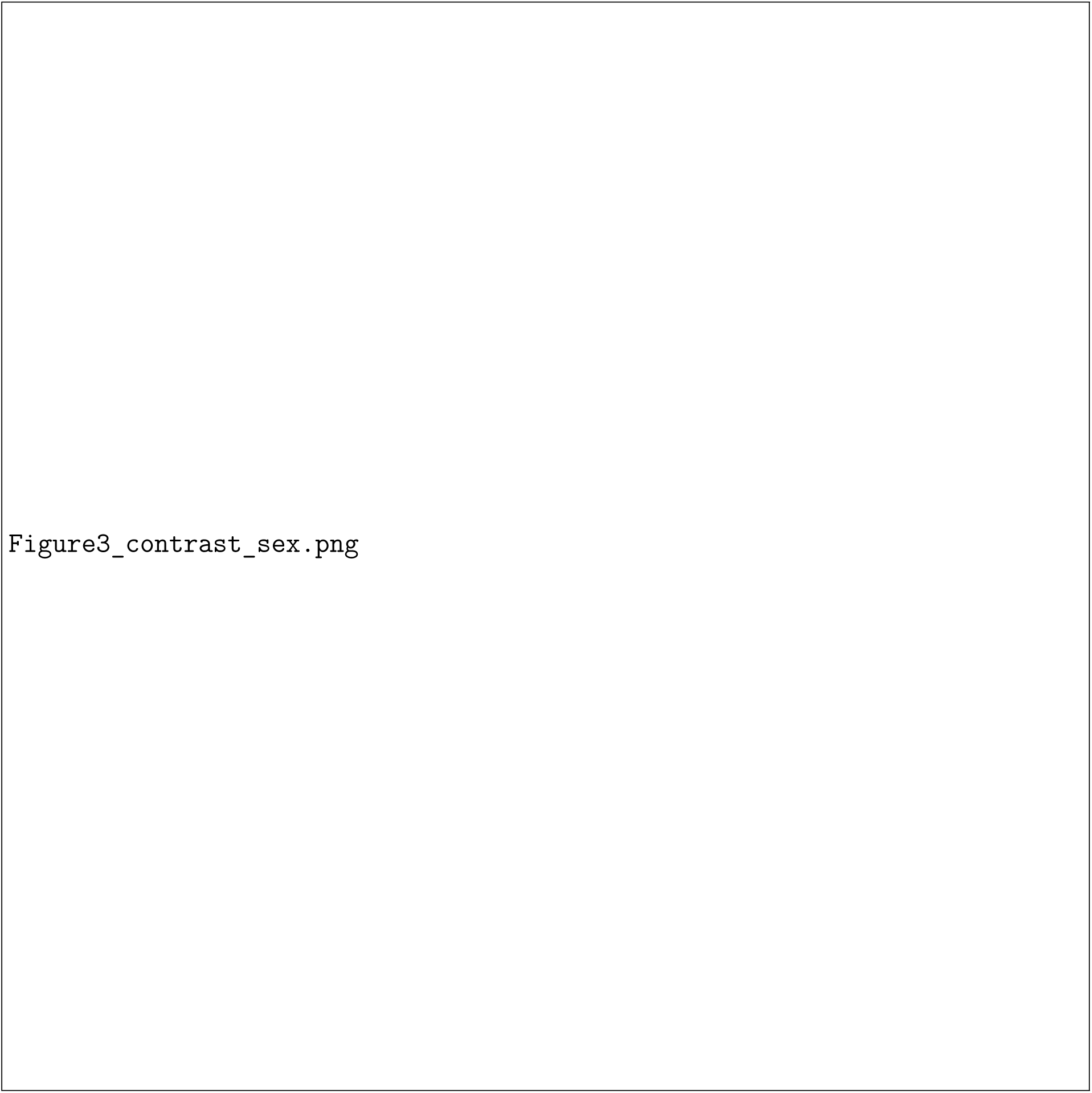
Omics analysis results for Alzheimer’s disease comparing disease and healthy conditions. (A) Volcano plot of differentially expressed genes. (B) Gene correlation matrix of the top 20 differentially expressed genes. (C) Violin plot of the first principal component distribution. (D) Principal component analysis with PERMANOVA significance testing. (E) Pathway enrichment analysis results. (F) Cell count distributions across major brain cell types stratified by biological sex. (G) Sex-stratified differential expression scatter plots comparing log2 fold changes in female versus male cohorts across GABAergic neurons, glutamatergic neurons, and oligodendrocytes.

**Fig. 4.**
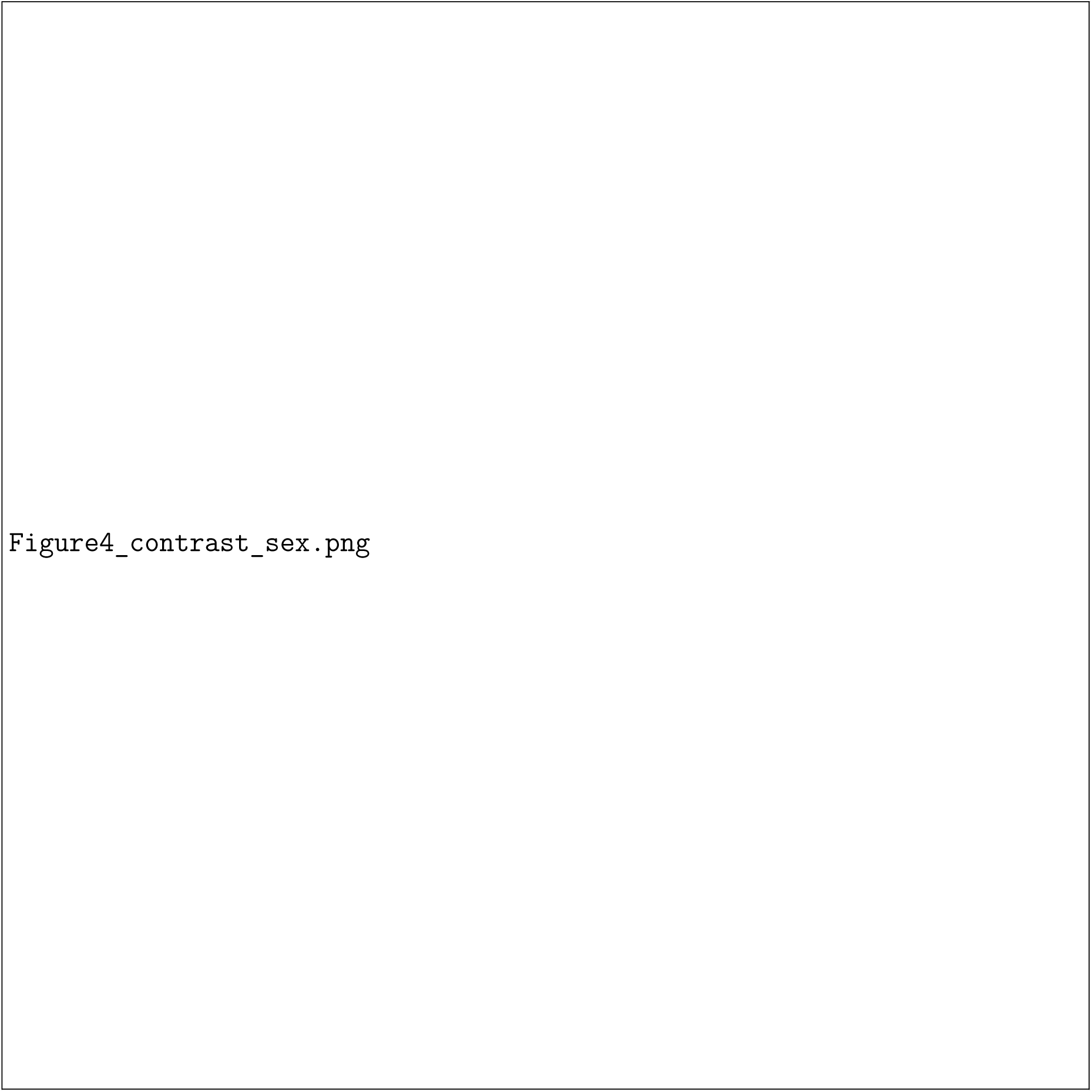
Omics analysis results for lung adenocarcinoma comparing cancer and normal conditions. (A) Volcano plot of differentially expressed genes. (B) Gene correlation matrix of the top 20 differentially expressed genes. (C) Violin plot of the first principal component distribution. (D) Principal component analysis with PERMANOVA significance testing. (E) Pathway enrichment analysis results. (F) Cell count distributions across major cell populations stratified by biological sex. (G) Sex-specific KEGG pathway enrichment bubble chart detailing female-enriched and male-enriched functional pathways across distinct cell types. (H) Sex-stratified differential expression scatter plots comparing log2 fold changes in female versus male cohorts across B cells, epithelial cells, and malignant cells.

At the pathway level, these transcriptional changes were organized into two major modules. The first was an upregulated immune/stress-response module, including cytokine-associated signaling and MAPK-related pathways. The second was a downregulated cellular maintenance module, including oxidative phosphorylation, mitochondrial complex I activity, ribosome-associated functions, and endoplasmic reticulum-linked protein processing. This overall pattern is concordant with established views of AD as a multi-system disorder involving glial activation, metabolic dysfunction, and proteostatic failure rather than a purely amyloid-centric process [5].

Among the upregulated genes, IL3RA showed the strongest alignment with prior AD literature. The report identified IL3RA upregulation together with enrichment of immune signaling pathways and knowledge-graph links to IL-3/GM-CSF and JAK-STAT signaling. This result is consistent with evidence that astrocyte-derived IL-3 programs microglia in AD and that microglia upregulate IL-3R*α* in the vicinity of *Aβ* pathology, supporting a role for this axis in disease-associated glial responses [28].

Taken together, these findings support the interpretation that the AD dataset captures a bona fide neuroimmune signal involving the IL-3/IL3RA axis. Accordingly, the present data support IL3RA as a robust disease-associated marker and a plausible mechanistic entry point, while future work should test whether chronic IL3RA signaling induces stable chromatin remodeling or prolonged inflammatory state transitions in microglia.

### Sex-stratified analysis uncovers cell-type-specific and sex-biased transcriptomic dysregulation in alzheimer disease

To further characterize the molecular heterogeneity within the Alzheimer disease cohort, OmniCellAgent performs a sex-stratified comparison across major brain cell types. Cell type composition analysis initially reveals sex-dependent differences in cell abundance, where male subjects contribute more GABAergic neurons and oligodendrocyte precursor cells, while female subjects exhibit higher proportions of fibroblasts and endothelial cells (Figure 3F). Sex-stratified log2 fold change scatter plots across three cell types demonstrate distinct gene dysregulation patterns. Oligodendrocytes are primarily characterized by male-specific upregulation involving 1943 genes, whereas glutamatergic neurons are predominantly driven by female-specific upregulation comprising 3460 genes. Male-specific upregulation is also prominent in GABAergic neurons with 1582 genes. Notable outlier genes exhibit distinct sex biases, with *MALAT1* showing concordant upregulation across both sexes, *MT-ND3* displaying male-specific upregulation, and *SNHG14* demonstrating strong female-specific upregulation. These results indicate that while a shared core of Alzheimer disease transcriptional response exists between sexes, a substantial proportion of the dysregulation is highly sex-specific and varies extensively across different cell types (Figure 3H).

### Pancreatic ductal adenocarcinoma exhibits transcriptomic rewiring toward RNA processing, nutrient scavenging, and microenvironmental adaptation

Differential expression analysis of the revised pancreatic ductal adenocarcinoma (PDAC) cohort identified a broad transcriptional program more consistent with adaptive remodeling than with isolated activation of canonical kinase pathways. In the report, BDP1, LENG8, and AP2A2 were prioritized as leading gene-level signals, and the concluding synthesis proposed that PDAC survival in this cohort may be supported by non-canonical RNA processing and aggressive nutrient-scavenging behavior. The revised report also emphasizes that this interpretation arises from a cohort of 828 samples and therefore requires explicit handling of tissue heterogeneity before strong mechanistic conclusions are drawn (Supplementary Information).

At the pathway level, PDAC is well known to survive in a nutrient-poor microenvironment by increasing extracellular nutrient uptake and scavenging pathways, including macropinocytosis and related endocytic processes. Recent work and reviews continue to support nutrient scavenging as a central feature of PDAC fitness under metabolic stress [39]. In that context, the report’s interpretation of AP2A2 as part of an uptake- and trafficking-oriented state is reasonable as a systems-level result, even though AP2A2 itself should still be treated as a prioritized candidate rather than an established PDAC driver.

The same principle applies to the RNA-processing axis. BDP1 is a core component of the TFIIIB complex required for RNA polymerase III transcription initiation, making it a credible anchor for altered transcriptional control [15]. LENG8 has emerging evidence as a regulator of mRNP quality control and nuclear RNA handling, supporting the report’s decision to connect it to post-transcriptional regulation, although the specific “viral mimicry” mechanism remains a hypothesis rather than a result [7]. Accordingly, the strongest interpretation is that the revised PDAC transcriptome converges on a plausible axis of RNA biogenesis, RNA handling, and stress-adaptive growth.

The principal analytical challenge in this single-cell setting is to resolve which malignant, stromal, or hypoxia-associated cell states express these genes, whether these programs co-occur within the same cells, and whether they mark aggressive or therapy-resistant subpopulations. Accordingly, the gene-specific interpretations are best treated as future directions rather than established conclusions. In particular, the proposed mapping of *AP2A2* and *HCG17* to hypoxic or stromal niches by spatial transcriptomics, together with experimental interrogation of the *LENG8/BDP1* axis, represents a logical next step for validating these hypotheses. More broadly, integration with the canonical PDAC genomic landscape remains necessary, because recurrent alterations in *KRAS*, *TP53*, *CDKN2A*, and *SMAD4* continue to define the core biology of PDAC and should inform interpretation of any transcriptomic state model [45].

The final case narrative can also be distilled into a synthesized network plot that links prioritized claims to their supporting omics context and translational hypotheses. For the PDAC case, this layer is particularly interpretable because the major report modules are also visible in the DEG and enrichment outputs, including transcriptional addiction, mitochondrial suppression, FBXO42–PP2A-associated chemoresistance, and nutrient-scavenging/endocytic behavior. As shown in Figure 5, the network plot is most appropriately read as a reporter-derived integration of report claims, bio evidence, and graph-level contextualization rather than as a direct molecular-interaction diagram. This representation provides a compact way to inspect which modules are strongly aligned across the report, the omics evidence, and the final synthesized graph.

**Fig. 5.**
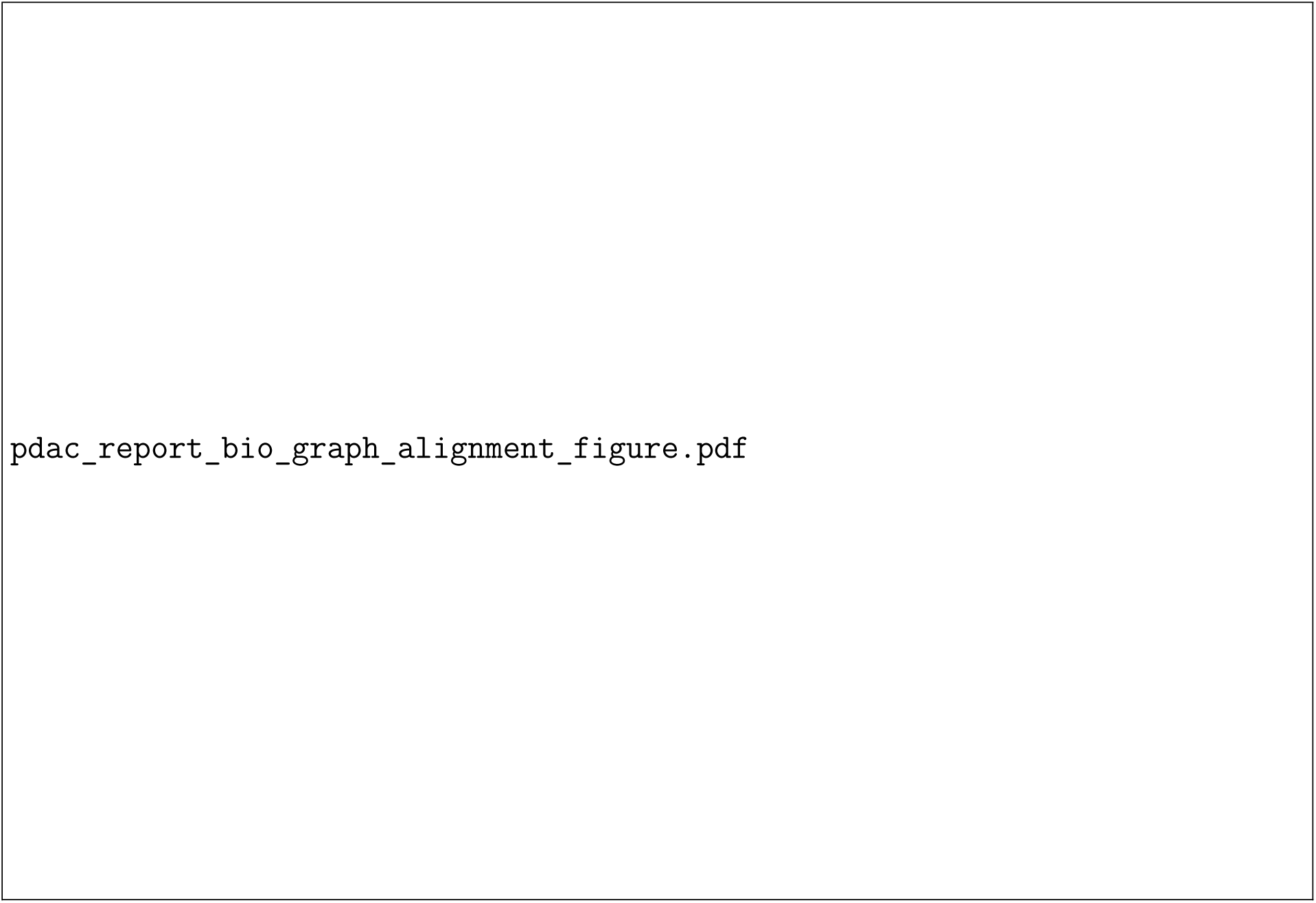
Illustrative report–bio–graph alignment for the PDAC case. The left column summarizes four core claims extracted from the revised Reporter-generated narrative, the middle column shows the corresponding bio-evidence bridges from differential expression, enrichment analysis, and pathway plots, and the right column highlights the matched modules within the final reporter-derived network plot. This example shows how the network layer functions as a compact synthesis artifact that links structured omics evidence to higher-level mechanistic and translational interpretation, rather than as a raw knowledge-graph or direct molecular-interaction output.

### Lung adenocarcinoma shows convergent mitochondrial, structural, and epigenetic dysregulation, with key gene-level signals requiring artifact-aware validation

In the revised LUAD report, differential expression analysis of 1,918 samples identified a striking transcriptomic pattern marked by strong upregulation of mitochondrial pseudogenes and downregulation of genes linked to mitochondrial structure, chromatin regulation, and cytoskeletal organization, including MAP4K3-DT, MT-ATP8, ZZZ3, TAFAZZIN, and SEPTIN2. The report further summarizes the affected pathways as mitochondrial bioenergetics, MAPK/mTOR-related signaling, and chromatin remodeling, supporting a systems-level LUAD phenotype centered on metabolic and transcriptional reprogramming (Figure 4).

At the pathway level, the mitochondrial and epigenetic interpretations are biologically plausible. Tafazzin is the phospholipid transacylase that remodels cardiolipin, a key inner-mitochondrial-membrane lipid required for oxidative phosphorylation and mitochondrial organization, making the report’s TAFAZZIN/MT-ATP8–linked mitochondrial vulnerability directionally reasonable [33]. Likewise, ZZZ3 is a histone H3 reader within the ATAC histone acetyltransferase complex, which is required for maintenance of histone acetylation and active transcription, while the broader ATAC complex, particularly YEATS2, has been implicated in NSCLC tumorigenesis [29].

The LUAD analysis is strongest where it explicitly states its technical limitations. It warns that the mitochondrial pseudogene signal may reflect read-mapping artifact and recommends re-alignment with strict unique-mapping filters before treating that signature as biological. That caution remains appropriate in single-cell data, where sparse counts and alignment ambiguity can distort interpretation of mitochondrial features. Likewise, the inconsistent behavior of *SEPTIN2* relative to prior NSCLC literature should be framed as a question of cluster-specific expression and malignant-state specificity. For that reason, the most novel single-gene claims, especially for *MAP4K3-DT* and *SEPTIN2*, are best presented as candidate biomarkers and future perturbation targets rather than established drivers.

### Sex-stratified analysis reveals divergent immune landscapes and pervasive male-specific transcriptional suppression in lung adenocarcinoma

To further investigate sexual dimorphism within the lung adenocarcinoma microenvironment, OmniCellAgent conducts a sex-stratified analysis across major cell populations. Cell type composition analysis initially reveals distinct sex-dependent differences in cell abundance. Female subjects exhibit a substantially higher proportion of T cells, macrophages, endothelial cells, and dendritic cells, whereas male subjects contribute a greater number of B cells, fibroblasts, and malignant cells. KEGG pathway enrichment analysis further elucidates divergent biological functional biases between sexes within the tumor microenvironment. Female-enriched pathways predominantly concentrate on classical immune surveillance and antigen presentation functions, with significant enrichment in pathways related to antigen processing, allograft rejection, and autoimmunity within epithelial cells and macrophages. Conversely, male-enriched pathways demonstrate significant metabolic reprogramming and broad inflammatory response characteristics. In male epithelial cells specifically, ribosome and oxidative phosphorylation pathways are highly active, while various male immune cells including macrophages, mast cells, plasma cells, and B cells show broad enrichment for proteasome, NF-kappa B, Toll-like receptor, and chemokine signaling pathways.

At the gene expression level, sex-stratified log2 fold change scatter plots of three key cell types corroborate this profound sex-specific transcriptomic remodeling. In malignant cells, both sexes exhibit large-scale gene downregulation comprising 4720 genes in males and 4716 genes in females, yet the number of female-specific upregulated genes reaching 1323 notably exceeds the 774 observed in males. However, within the non-malignant compartments, the dysregulation pattern presents an overwhelming male-specific transcriptional repression. Epithelial cells contain 3972 male-specific downregulated genes, and B cells display an extreme male-specific downregulation involving 12662 genes, with negligible female-specific changes. Furthermore, specific outlier genes indicate potential mechanistic divergences, such as the ribosomal protein gene *RPS9* demonstrating concordant upregulation across both sexes in epithelial cells, and the mitochondrial gene *MTCYBP19* showing significant concordant upregulation in B cells. These results suggest that transcriptomic dysregulation in lung adenocarcinoma is highly cell-type specific and is primarily driven by massive male-specific gene repression within immune and microenvironmental stromal cells.

## DISCUSSION

Real-world biomedical scientific discovery operates as an iterative lifecycle integrating four core pillars: the targeted identification and analysis of question-specific omics datasets; the context-aware interpretation of molecular data using rich biomedical prior knowledge database; comprehensive literature reviews; and the subjective creativity and expert intuition of human scientists. Together, these pillars drive robust evidence synthesis and novel hypothesis generation. The recently reported AI agents support automated omics analysis and literature review, which typically require users to predefine and curate disease-specific datasets, which is a process that remains challenging and time-consuming.

In omic-driven biomedical research, identifying and analyzing research question-relevant omics datasets, and interpreting analytical results using prior knowledge from databases and literature review, together with scientist-specific research taste, creativity and intuition, are essential for generating novel scientific hypotheses and discoveries. Several agentic AI frameworks have recently been developed to automate omics analysis and literature review, facilitating omics-driven studies. However, critical challenges remain. Identifying and curating disease-control paired omics datasets from heterogeneous public repositories is still labor-intensive and requires substantial domain expertise. Interpreting top-ranked targets further demands integration with complex, multi-source biological knowledge. In addition, current systems have limited ability to incorporate expert-specific research perspectives during data interpretation.

In this exploratory study, we present OmniCellAgent, a multi-agent AI framework that integrates large-scale single-cell omics data, biomedical prior knowledge and literature-driven reasoning to support omics-driven scientific discovery. By unifying dataset retrieval, analysis, knowledge grounding and hypothesis generation within a single system, OmniCellAgent addresses a key bottleneck in precision medicine: the fragmentation of data, tools and expertise required for hypothesis generation. Across multiple disease settings, the framework recovers biologically meaningful signals, prioritizes candidate targets and produces structured, evidence-supported hypotheses. It coordinates otherwise isolated AI agents and tools into workflows that resemble practical scientific analysis. Through a planner-executor-reporter architecture with structured state sharing, OmniCellAgent enables multi-step reasoning while preserving critical intermediate outputs such as gene lists and pathways. This separation of structured data flow from language-based reasoning improves robustness and interpretability.

In the case studies, OmniCellAgent consistently identifies biologically plausible targets across disease contexts. For example, in Alzheimer’s disease, it recovers convergent immune and metabolic dysregulation, including IL3RA-associated neuroinflammatory pathways. The sex-stratified analysis within the lung adenocarcinoma case study further illustrates the translational value of the layered outputs of OmniCellAgent. By explicitly mapping the cell-type abundance baseline before differential testing, the framework alerts researchers to potential proportion-driven biases, a confounding factor that is frequently overlooked in heterogeneous tumor microenvironment analyses. The pathway-level polarization map translates sex-biased immune signatures, including female-enriched antigen processing and presentation activity in epithelial cells and macrophages alongside male-enriched metabolic reprogramming, into cell-type-resolved hypotheses that could inform the development of sex-stratified preclinical models and guide future therapeutic investigations. The identification of profound sex-specific transcriptional divergences, such as the massive male-biased gene repression in B cells and the emergence of discordant genes, albeit limited in number, with opposing regulatory directions within malignant cells, suggests potential biological vulnerabilities where uniform therapeutic modulation might have divergent effects across patient populations These findings position such genes as cautionary markers for target selection and priority candidates for sex-specific biomarker development. For the other case studies, it identifies supportive evidence of intersting targets, pathways and drugs. These results extend beyond single-model or single-pipeline outputs by synthesizing information across genomics, knowledge graphs and the scientific literature to identify non-obvious, systems-level connections. They demonstrate the system’s ability not only to detect omics signals but also to generate context-aware interpretations grounded in data, prior knowledge and literature.

Furthermore, for open-ended questions, the agent does more than list facts and sources from tools. It can potentially highlight inconsistent findings within the retrieved literature, contrast varying scientific perspectives, and provide users with synthesized summaries to support their own expert judgment, new ideas and hypothesis refinement. In this way, OmniCellAgent is not merely a closed-form question-answering system, but a platform that augments researchers working on open scientific questions. The agent is well suited to exhaustive, data-intensive tasks such as identifying patterns and potential connections, freeing human researchers to focus on experimental design, creative problem-solving and interpreting biological meaning within a broader scientific context. By streamlining complex data analysis, OmniCellAgent empowers biomedical researchers and biological domain experts to engage more directly with omics data without requiring extensive independent coding pipelines.

In summary, OmniCellAgent illustrates the potential of multi-agent AI systems as AI co-scientists capable of integrating omics data, biomedical prior knowledge, literature review and question reasoning to facilitate and accelerate evidence collection and hypothesis generation in the AI-human collaboration manner. By reducing barriers to omics-driven analysis and enabling structured, multi-step discovery, this work highlights a shift toward AI systems that function as integrative engines for scientific workflows.

Although highly promising for accelerating discovery, multi-agent AI for science is currently in a preliminary stage with inherent limitations. There are still some limitations in this exploratory study. For example, the future work will focus on scaling to additional single-cell omics modalities beyond scRNAseq (like paired scATACseq and scRNAseq), incorporating omics-analysis foundation models (FMs) pre-trained on large-scale single-cell omics data, expanding reliable and effective multimodal data analysis and evidence-synthesis functions for novel and testable hypothesis generation, collaborating with biomedical experts for hypothesis validation across disease studies, and enabling closed-loop experimental validation to further advance agentic AI frameworks for omics-driven precision medicine.

## METHODOLOGY

### Methodology Overview

OmniCellAgent is implemented as a LangGraph state machine in which planner, executor, replanner and reporter nodes operate over a shared `AgentState`. The planner converts each query into a structured `ResearchPlan` of typed subtasks; the executor dispatches subtasks and stores their outputs; the replanner retries or revises failed steps; and the reporter produces a citation-linked synthesis. The shared state stores the query, plan, candidate genes, pathways, disease and cell-type context, literature identifiers, per-agent outputs and process log, decoupling agent coordination from raw conversational context.

The Omic Data Agent maps disease, tissue and cell-type terms from the query to OmniCellTOSG [62], retrieves matched disease-control expression matrices, and performs differential expression analysis with a parallel Wilcoxon rank-sum test followed by GO and KEGG enrichment. Unless otherwise specified, genes with FDR < 0.05 were treated as significant. For each case study, the agent generated PCA scatterplots, PERMANOVA tests on Bray–Curtis distance, Pearson correlation summaries of the top differentially expressed genes, and dot or lollipop enrichment plots, then wrote these outputs to state for downstream agents (Figure 6; full workflow details are provided in the Supplementary Information).

**Fig. 6.**
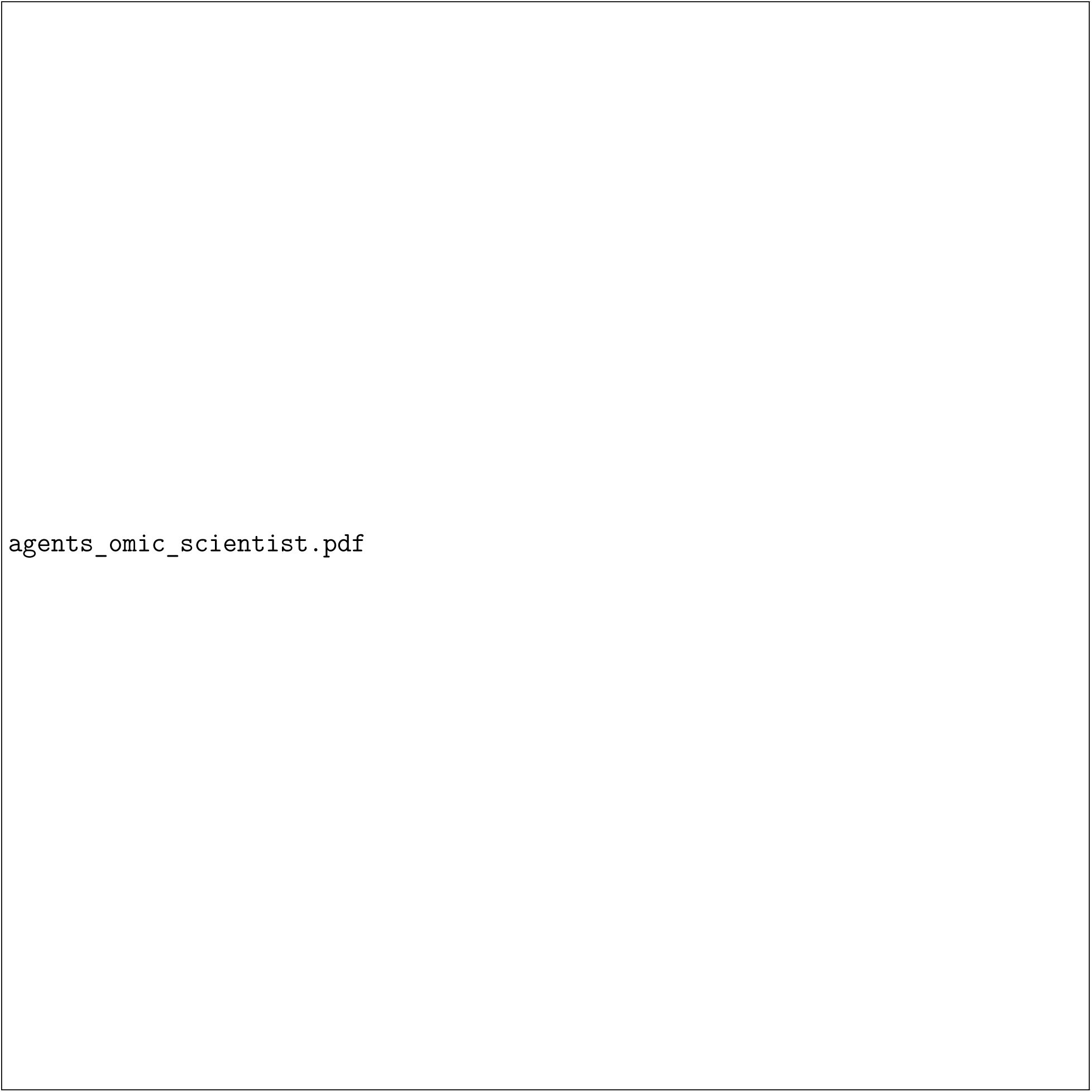
Omic Data Agent and Scientist Expert Agent workflows. The Omic Data Agent maps query entities to OmniCellTOSG, performs differential-expression and enrichment analyses, and writes ranked genes, pathways and plots to the shared `AgentState`. The Scientist Expert Agent consumes accumulated omics, knowledge-graph and literature evidence, routes the question to domain experts, and synthesizes ranked mechanistic hypotheses and validation ideas.

The BioMarkerKG Agent queries a PrimeKG-derived biomedical knowledge graph built from STaRK-Prime [58], using graph retrieval and a graph language model to connect candidate genes with diseases, pathways, drugs and protein interactions. The PubMed Agent groups candidate genes into focused queries, retrieves article metadata and available full text through the NCBI E-utilities API, and returns DOI/PMID-linked summaries. These evidence-retrieval agents are summarized in Figure 7; detailed workflow diagrams are provided in the Supplementary Information. The Google Agent adds complementary web-scale evidence for translational or recently emerging context.

**Fig. 7.**
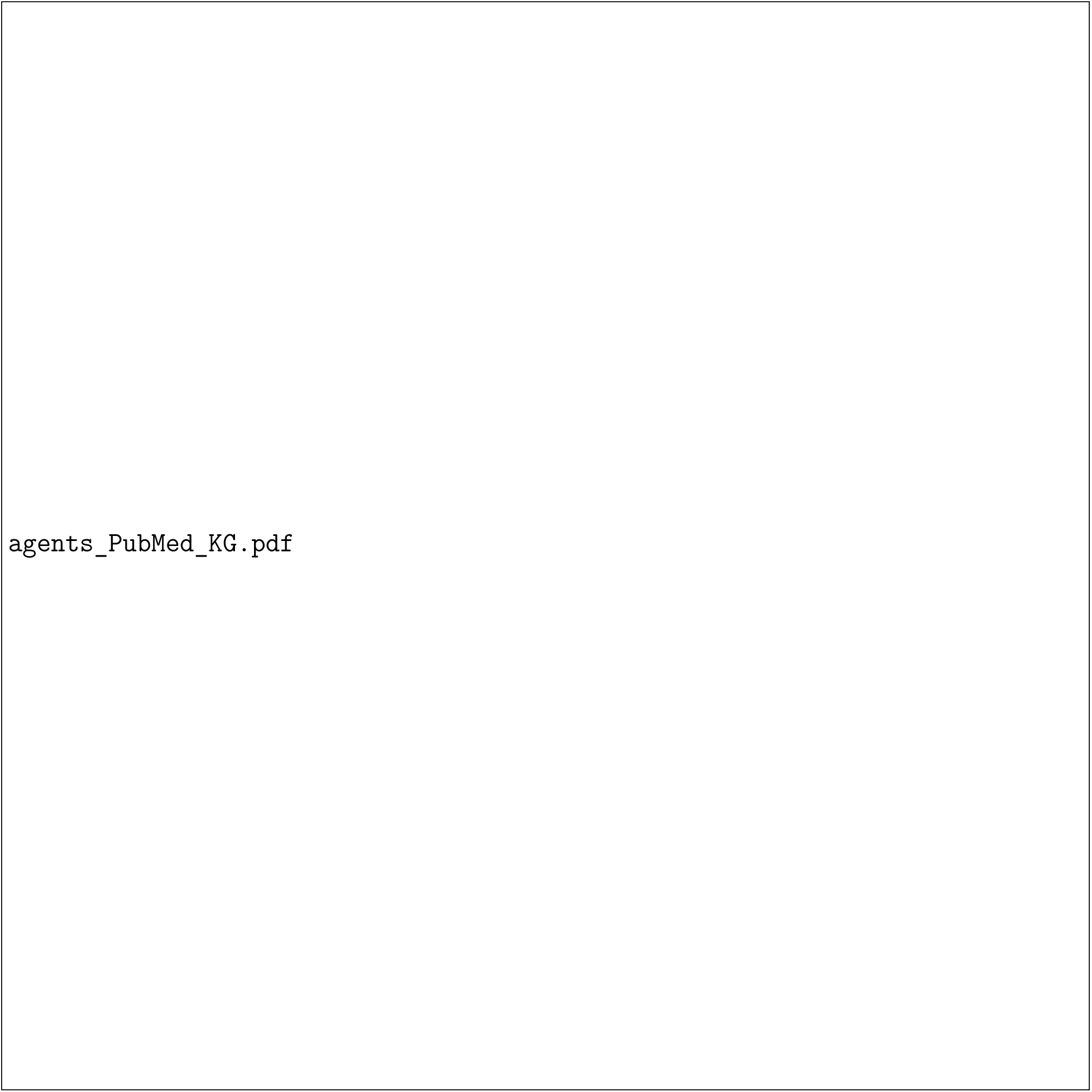
PubMed Agent and BioMarkerKG Agent workflows. The PubMed Agent builds focused literature queries from candidate genes and prior agent outputs, retrieves DOI/PMID-linked evidence, and returns concise summaries for downstream synthesis. The BioMarkerKG Agent queries a PrimeKG-derived biomedical graph to connect candidate genes with pathways, diseases, drugs and protein interactions, using graph retrieval and graph-language-model reasoning to produce grounded biomarker evidence.

The Scientist Expert Agent routes questions to domain-specific experts (Genomics, Neuroscience, Longevity/Biostatistics, Cancer) backed by pre-built LightRAG knowledge bases [17] of curated publication corpora. It synthesizes ranked mechanistic hypotheses with evidence-strength scores and proposed validation experiments. The workflow is summarized with the Omic Data Agent in Figure 6; additional workflow and expert knowledge-base details are provided in the Supplementary Information.

The Reporter Node integrates the task plan, agent outputs, shared data and generated figures into a structured report covering omics results, knowledge-graph context, literature-supported targets, pathway enrichment, ranked hypotheses, validation suggestions and references. A programmatic appendix preserves session metadata, the execution plan, process log and truncated agent outputs, allowing each hypothesis to be traced back to its upstream evidence. Detailed prompts, reports and per-agent diagrams are provided in the Supplementary Information.

### Methodology

The complexity of hybrid data sources and the necessity of multi-step reasoning, where each step depends on the previous ones, yield the requirement for a *multi-agent system*, which needs an appropriate and effective *orchestrator*. The Orchestrator is implemented as a state machine using *LangGraph*, designed to manage the lifecycle of a hypothesis-driven discovery workflow. It routes subsequent agents to complete their corresponding local sub-tasks, in service of the main goal of scientific research analysis with single-cell omics. Unlike linear execution models, the Orchestrator maintains a persistent *AgentState* that tracks the user query, execution plan, shared data context—such as gene lists and DOI references—and process logs across distinct phases. This graph-based architecture ensures that the system is not just reactive but follows a robust *Plan-Execute-Replan* cycle, allowing for adaptive handling of complex biomedical queries. The main manuscript illustrates this architecture design.

### Omic Data Agent

The OmicDataAgent is a specialized component designed to facilitate context-aware multi-omics analysis for biomedical research queries. Its primary functions include: (1) identifying and retrieving omic datasets relevant to a given biological or disease-related question from OmniCellTOSG [62], (2) performing robust statistical analyses and enrichment assessments, and (3) extracting key biological insights such as differentially expressed genes (DEGs) and significantly enriched signaling pathways and biological processes. OmniCellTOSG (Omni-Cell Text-Omic Signaling Graph) is a large-scale, graph-structured, AI-ready dataset that harmonizes single-cell transcriptomics data and the BioMedGraphica biological knowledge graph [60, 61] across diverse disease contexts, tissue types, and cell populations. It integrates curated annotations, metadata, and multi-omics features from public repositories such as CellxGene [37], BrainCellAtlas [6], and GEO [9], enabling comprehensive representation of disease-relevant cellular states and signaling activities. The graph structure of OmniCellTOSG encodes both molecular attributes (e.g., gene expression profiles, pathway activities) and biological relationships (e.g., signaling pathways and protein-protein interactions), allowing intelligent agents to reason over complex omic landscapes. Figure 8 illustrates the internal mechanism of the Omic Data Agent.

**Fig. 8.**
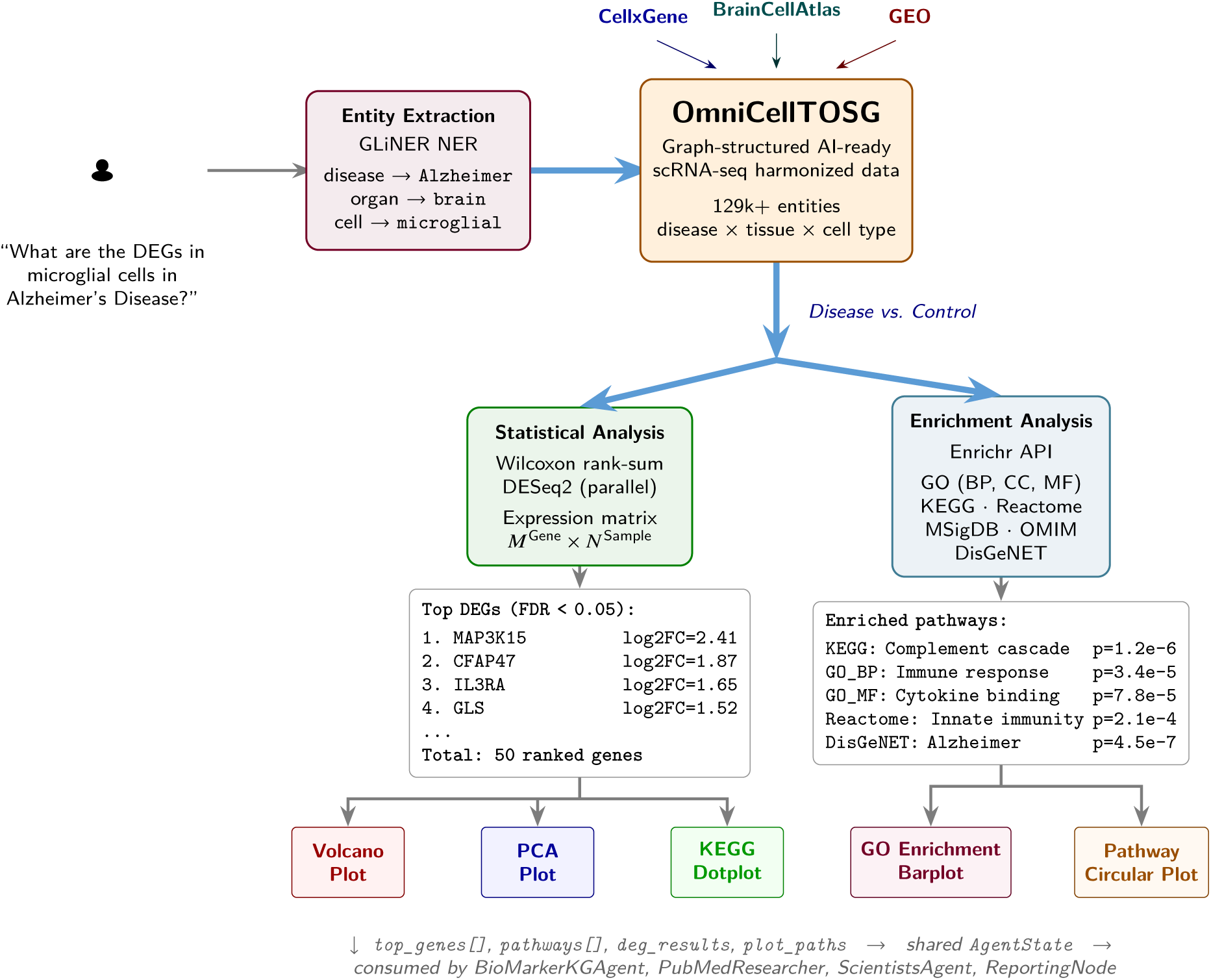
Internal mechanism of the Omic Data Agent. A user query is first parsed via GLiNER named-entity recognition to extract disease, organ, and cell-type entities, which are then used to query OmniCellTOSG—a graph-structured, harmonized scRNA-seq resource built from Cellx-Gene, BrainCellAtlas, and GEO. Retrieved disease-vs-control expression matrices undergo statistical testing (Wilcoxon rank-sum / DESeq2) to identify differentially expressed genes (DEGs), which are subsequently submitted to the Enrichr API for functional enrichment across GO, KEGG, Reactome, MSigDB, and DisGeNET. The pipeline outputs ranked DEG lists, enrichment tables, and a suite of visualization plots, all written to the shared `AgentState` for downstream agents.

**Fig. 9.**
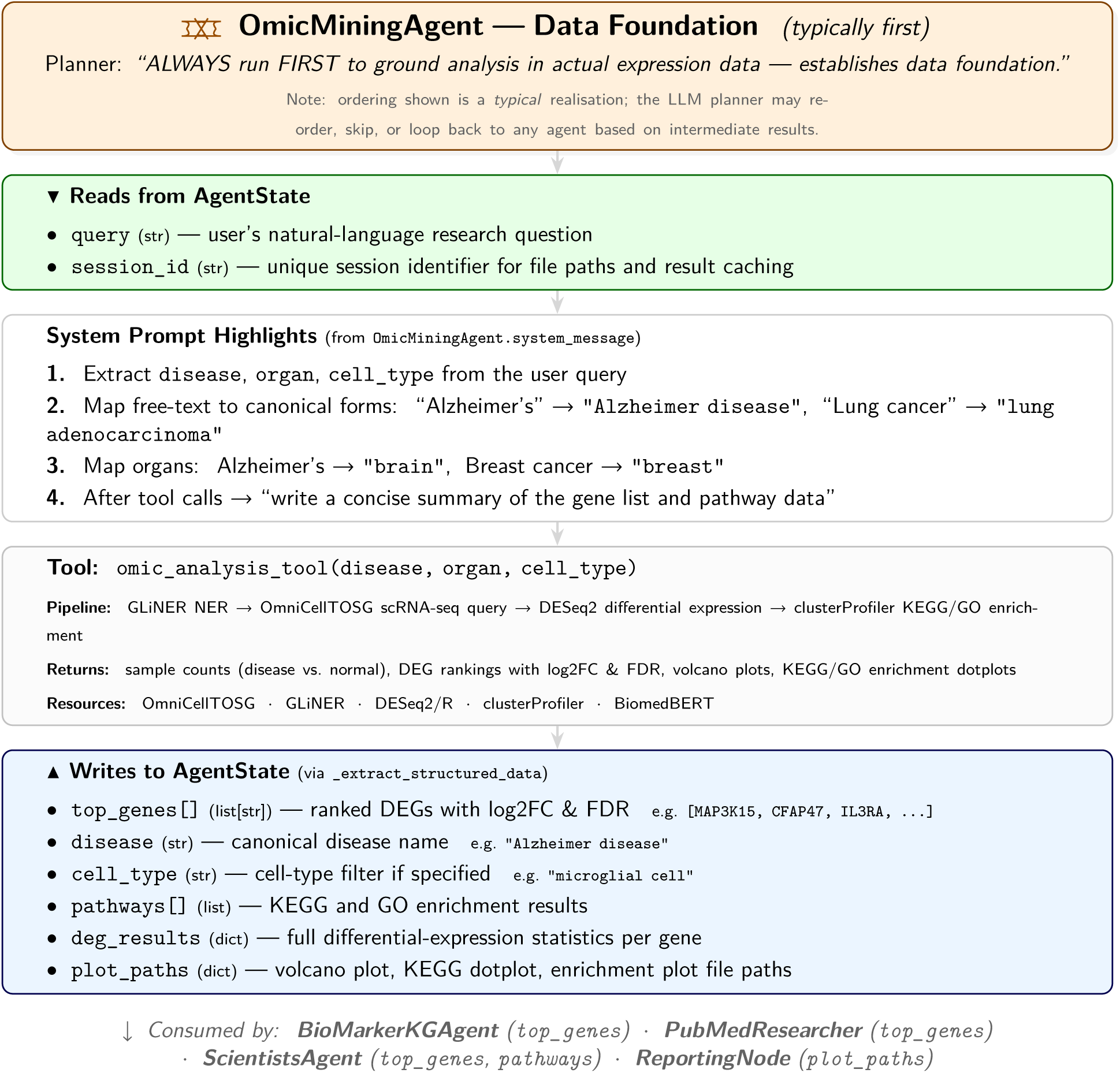
OmicMiningAgent. — workflow interface view. The ordering shown here reflects a *typical* plan generated by the LLM orchestrator; in practice the planner dynamically decides which agents to invoke, in what order, and whether to loop back after failures or surprising results. That said, the planning prompt strongly encourages running this agent first to “ground ALL hypotheses in actual omics data.” It reads only the user query and session identifier, maps them to canonical disease/organ/cell-type forms via its system prompt, and invokes a single tool (`omic_analysis_tool`) that chains NER, database retrieval, differential expression, and pathway enrichment. The six fields it writes to `AgentState`—most critically `top_genes[]`—become the shared anchor that every downstream agent reads.

**Fig. 10.**
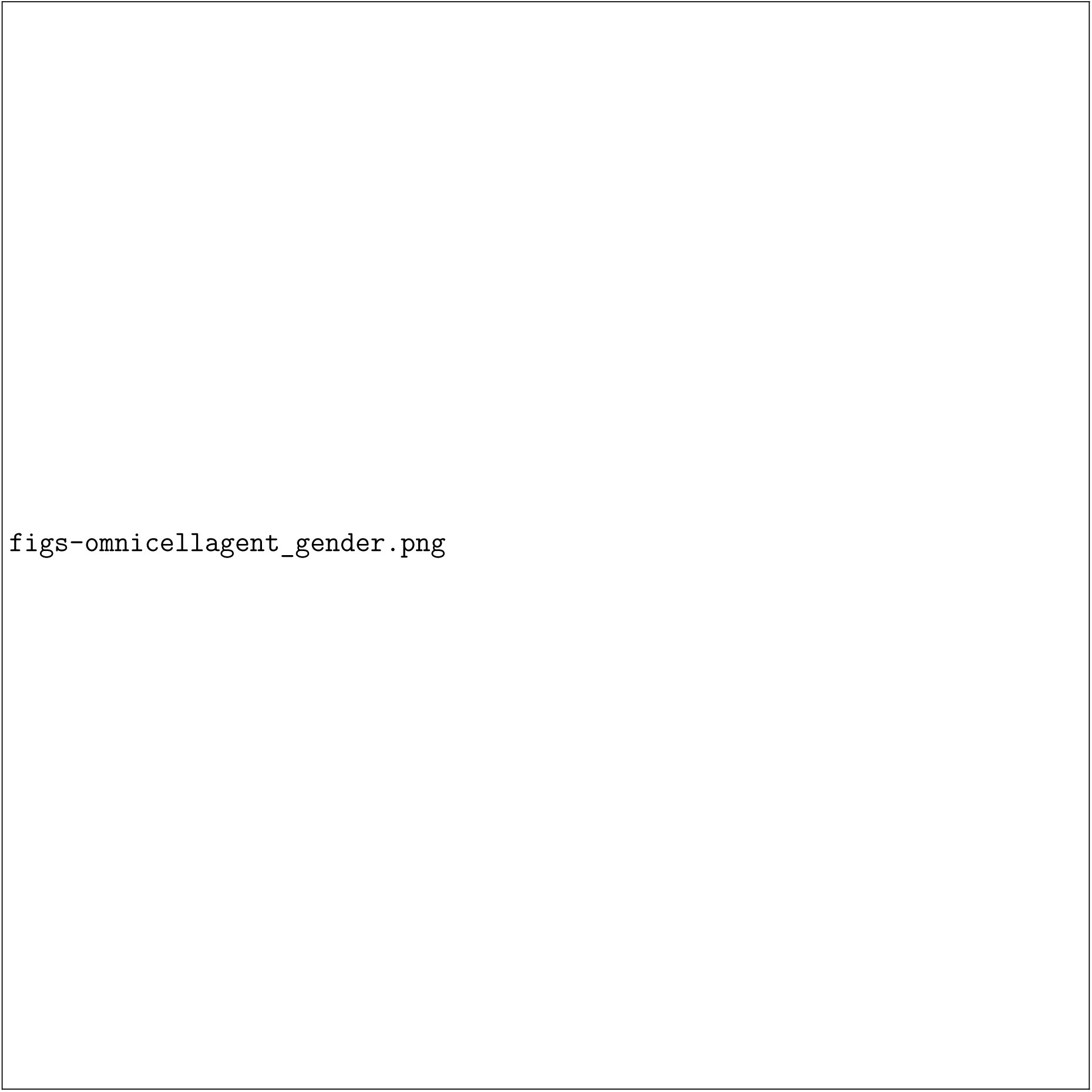
Omics analysis results for Alzheimer’s disease in microglia, comparing male versus female samples.

**Fig. 11.**
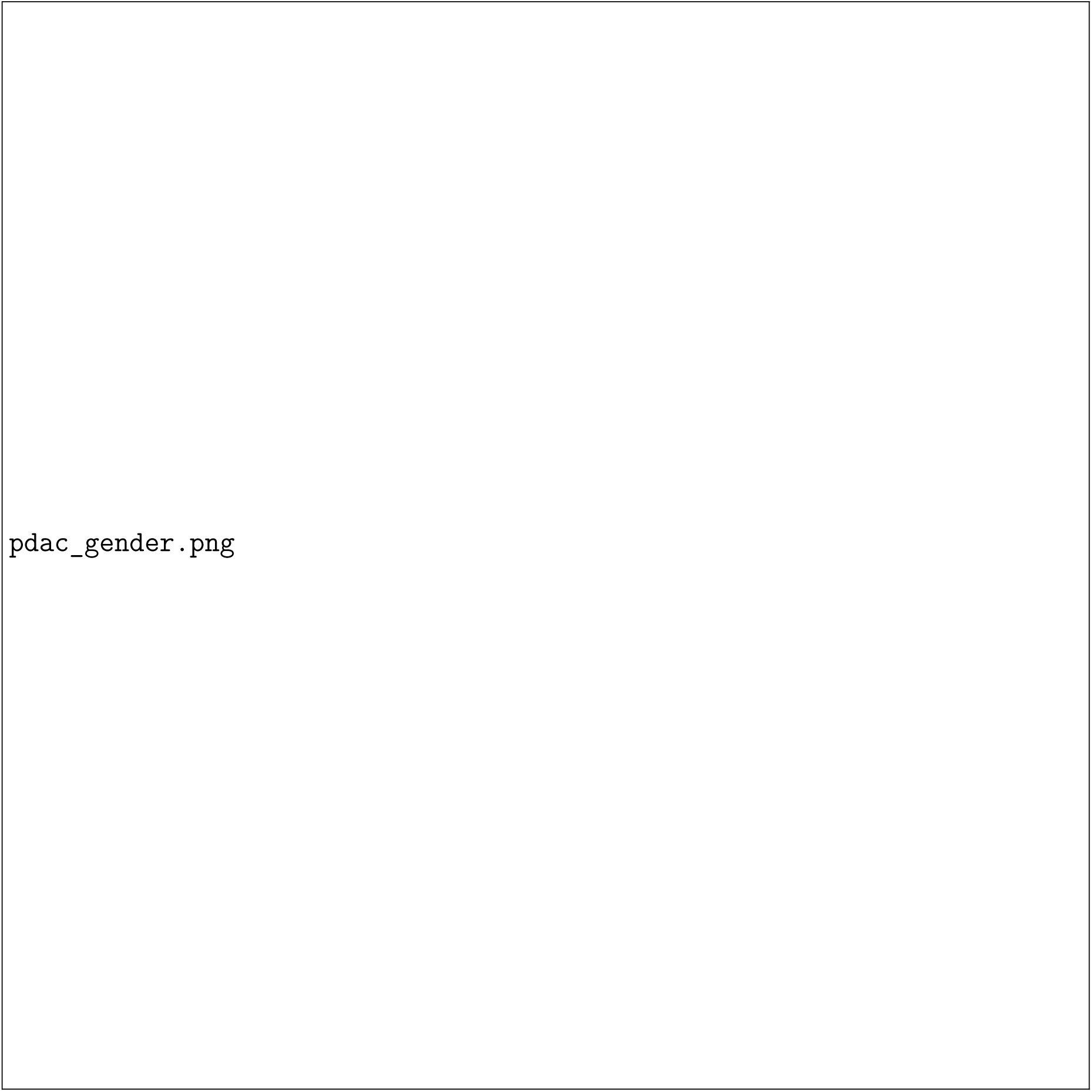
Omics analysis results for pancreatic ductal adenocarcinoma (PDAC), comparing male versus female samples.

When a user submits a natural language query, such as "What are the differentially expressed genes in microglial cells in Alzheimer’s Disease?", the Omic Data Agent interprets the prompt to extract key biological entities, including the disease context, relevant cell type, and intended analysis objective. It then retrieves matched disease and control meta-cell datasets from OmniCellTOSG, a harmonized resource of single-cell omics profiles curated across diverse biological and pathological conditions. The omic agent is able to filter relevant omic information from a list of tissues, cell types, diseases, patient ages, and sexes. We use Alzheimer’s disease as a case study to show the agent’s ability to analyze multi-omics data. Using statistical models such as the Wilcoxon rank-sum test (executed with parallel processing), the agent identifies differentially expressed genes (DEGs) within the specified context. It then outputs the expression matrices for disease vs. normal or female vs. male, which we use for downstream differential and pathway analysis. The extracted expression matrix *M*^*Gene*^ × *N*^*Sample*^ is 41,149 × 534 for disease vs. healthy and 41,149 × 1,296 for male vs. female. Principal component analysis (PCA) and Pearson correlation analyses were performed on the top 20 DE genes (scaled features). For each PC (PC1, PC2), group differences were summarized with a boxplot per subtype and compact letter displays derived from one-way ANOVA followed by Tukey’s HSD. PERMANOVA (Bray–Curtis distance on the same top-20-gene matrix) tested overall compositional differences between groups; the degrees of freedom, *r*^2^, and p-value were reported. A global Kruskal–Wallis test across groups was also computed and reported in the main disease-versus-control AD figure. Unless otherwise specified, genes with FDR < 0.05 were considered statistically significant.

These DEGs are subsequently analyzed through a functional enrichment workflow that programmatically interfaces with the Enrichr API. The agent submits gene lists and queries a wide array of curated databases, such as Gene Ontology (GO) [49], KEGG [21], Reactome, MSigDB, OMIM, and DisGeNET [34] — to uncover enriched biological pathways and disease associations. The analysis pipeline includes automated gene list submission, structured retrieval and storage of enrichment results across multiple databases, generation of summary reports distinguishing pathway- and disease-based enrichments, and export of top-ranked terms prioritized by adjusted p-values, along with annotated gene counts and pathway memberships.

Following enrichment analysis, the resulting gene set enrichment tables across Gene Ontology (GO) categories—Biological Process (BP), Cellular Component (CC), Molecular Function (MF)—as well as KEGG pathways and disease associations (from DisGeNET) were consolidated into a unified data structure for visualization. Each result table contained pathway terms, associated gene counts, and adjusted p-values. To support comparative interpretation, the top five most significant terms (ranked by adjusted p-value and gene count) were selected within each ontology category. The output consists of the AD disease-versus-control and LUAD figures shown in the main manuscript, plus the AD sex-stratified and PDAC figures included below, all produced using `ggplot2` [56].

### Scientist Expert Agent

The ScientistExpertAgent is designed to provide deep, specialized knowledge reflecting the cumulative work and historical perspective of a specific scientific expert. Its function is to serve as a focused repository of that individual’s contributions to their domain. The expert’s publications are routinely indexed and cached. This extensive corpus of scientific papers is processed and managed using the LightRAG framework [17]. Within LightRAG, documents undergo several processing steps: they are parsed and segmented into text chunks, and dense vector embeddings are generated for these chunks to enable efficient semantic similarity search across the large volume of text. LightRAG also constructs an internal knowledge graph from the ingested papers. This graph captures entities and relationships discussed within the expert’s work, offering a structured, multi-dimensional representation of their knowledge landscape beyond simple text retrieval.

Upon receiving a query, the Scientist Expert Agent retrieves relevant documents from its indexed corpus and generates a response based on the expert’s accumulated knowledge. This response is tailored to reflect the expert’s unique insights and perspectives, providing users with a specialized understanding of the topic at hand. Figure 14 illustrates the workflow of the Scientist Expert Agent. One important modularization is to have knowledge graph building modules separated from the agent call. The user first sets up the author names corresponding to each expert role, and a program builds the knowledge base that provides grounding to this scientist agent and stores it. This process is illustrated in Figure 12. In this way, one can build this knowledge base once and reuse it multiple times to save time and tokens.

**Fig. 12.**
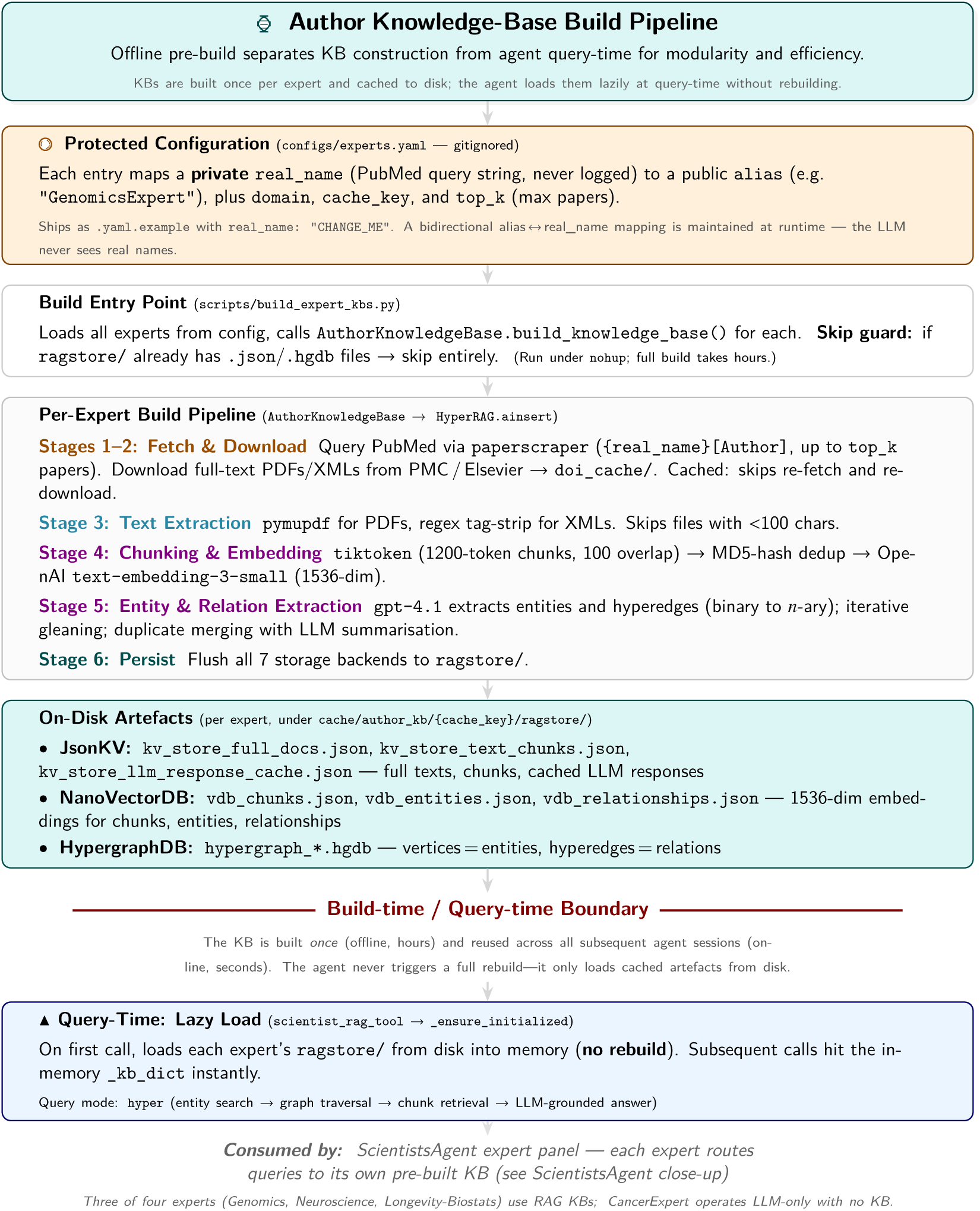
Author Knowledge-Base Build Pipeline. A git-ignored `experts.yaml` maps each expert alias to a scientist name used *only* for PubMed queries. The offline build (Stages 1–6) fetches papers, extracts text, chunks, embeds, and runs LLM entity/relation extraction. At query-time the pre-built KB loads from cache in seconds. This separation provides (i) *modularity* (experts updated independently), (ii) *efficiency* (build once, query many), and (iii) *privacy* (real names never reach the agent). See companion illustration for the HyperRAG retrieval architecture at query-time.

**Fig. 13.**
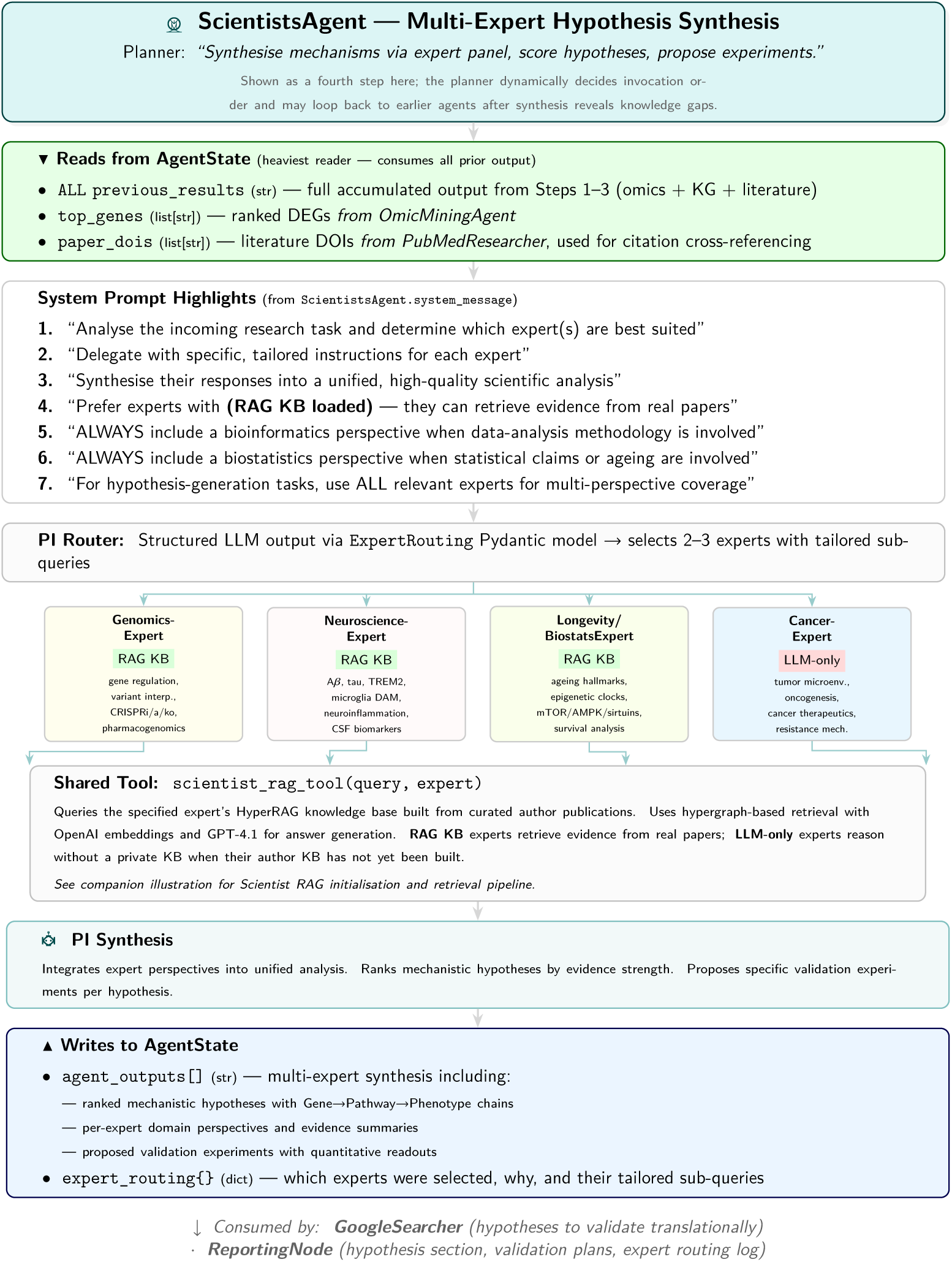
ScientistsAgent. — workflow interface view. The heaviest reader of `AgentState`, this compound agent consumes all prior omics, KG, and literature output. A PI-level router uses a structured `ExpertRouting` Pydantic model to select 2–3 of the four domain experts configured in `configs/experts.yaml` and dispatch tailored sub-queries. A fallback system decides whether to build or run the knowledge base or fallback to prompt only based on this configuration file. Genomics, Neuroscience, and Longevity/Biostatistics experts are backed by curated-author HyperRAG KBs (*RAG KB*); CancerExpert is configured but its KB is not yet built and is therefore routed *LLM-only*. The PI synthesis node integrates expert perspectives into ranked hypotheses with proposed validation experiments.

**Fig. 14.**
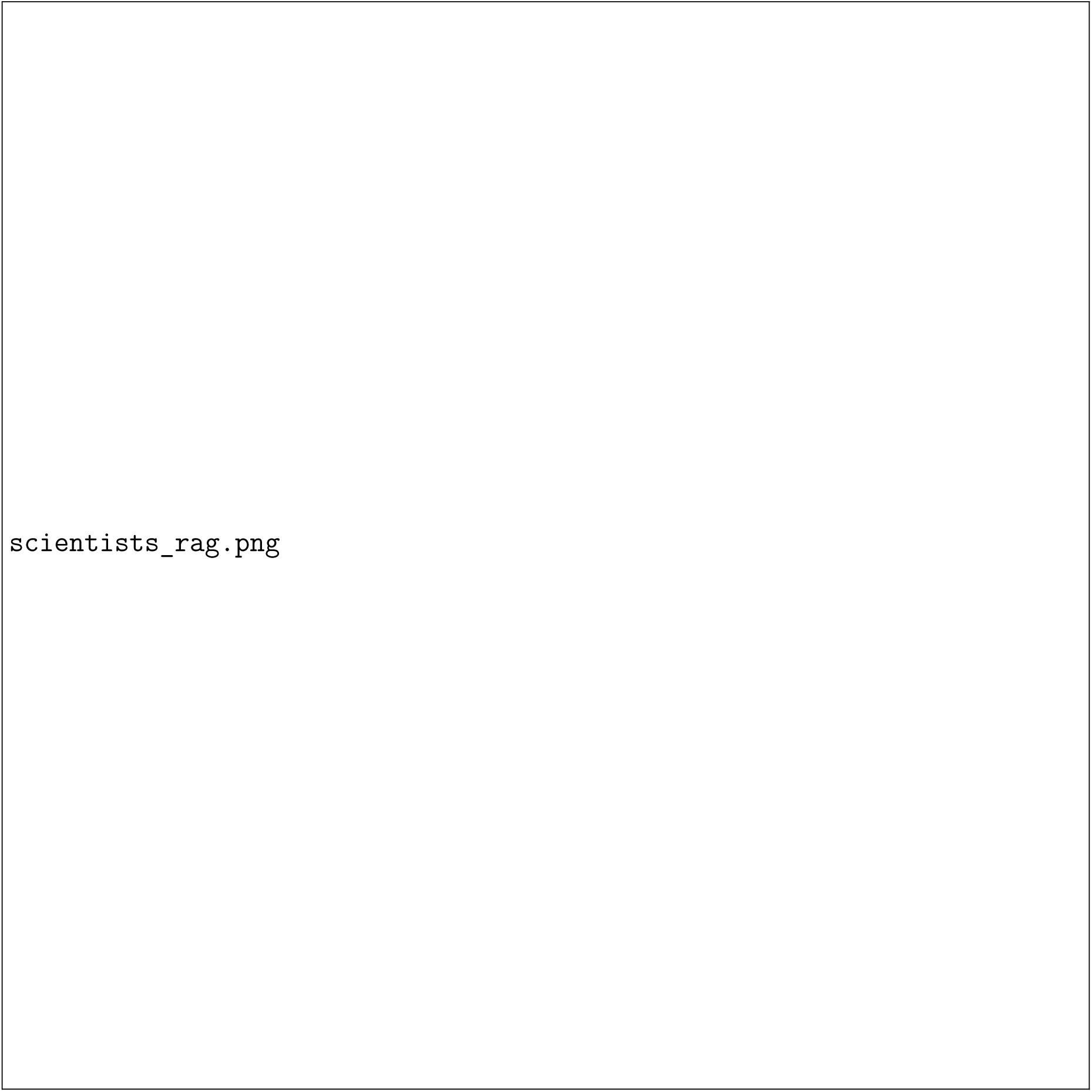
Overview of the Scientist Expert Agent workflow. The agent can be initialized with knowledge bases built from curated scientist publication corpora. At query time, retrieved entities, relations, and textual context are used to support expert-specific responses and downstream PI synthesis.

**Fig. 15.**
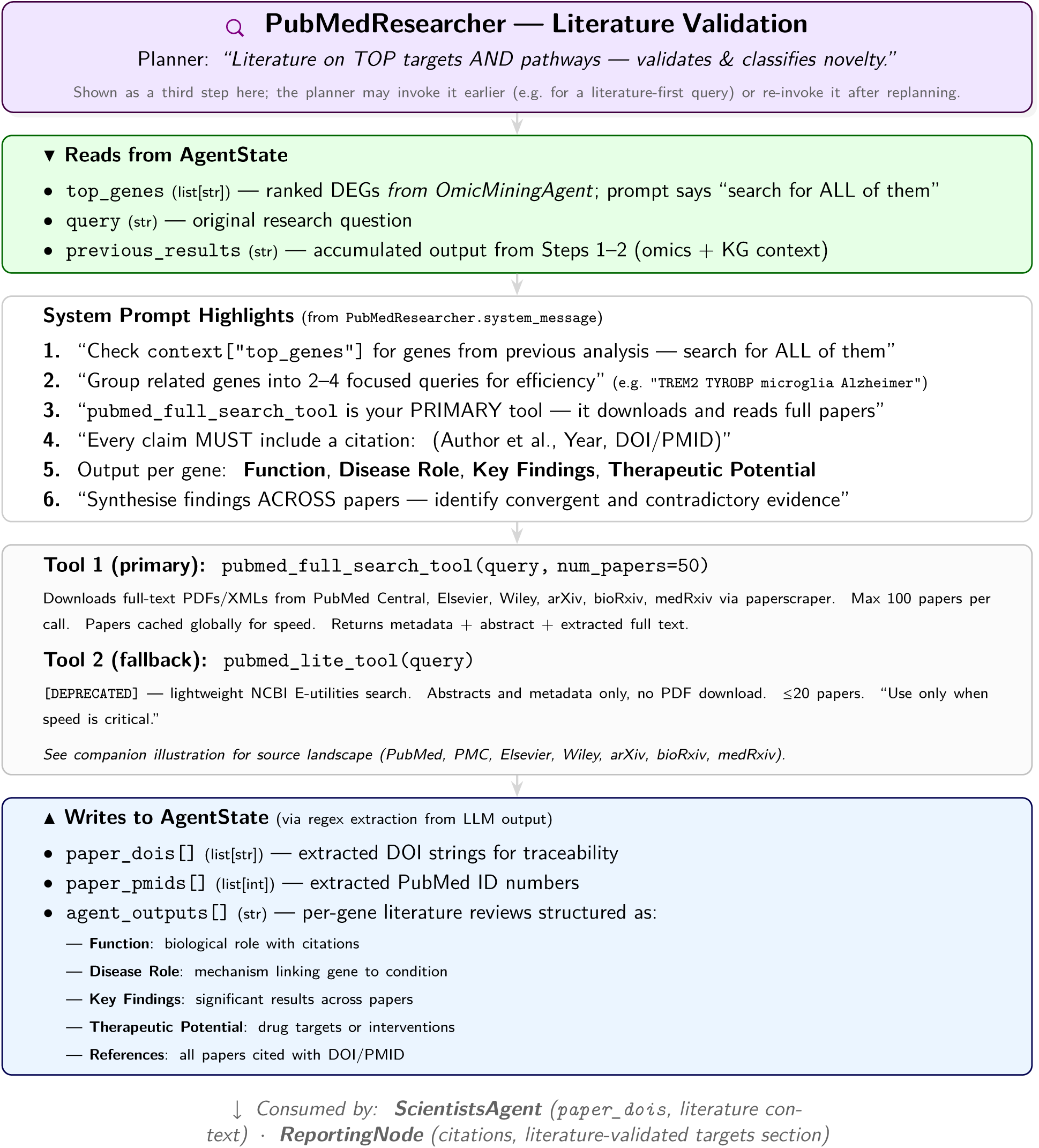
PubMedResearcher. — workflow interface view. Although shown here in the third position, the planner may invoke this agent at any point—even first for purely literature-driven queries—and may re-invoke it after replanning if earlier agents fail or surface unexpected targets. It provides the evidence layer. Its system prompt instructs it to read *all* genes from `top_genes` and group them into 2–4 focused queries for efficient retrieval. The primary tool downloads up to 100 full-text papers per call with global caching; a deprecated lite fallback is available for speed-critical lookups (see companion illustration for source landscape). Every claim is required to carry a citation. The structured per-gene output and extracted DOIs/PMIDs it writes to `AgentState` provide the literature grounding that ScientistsAgent uses for hypothesis novelty classification and that ReportingNode uses for the citation apparatus.

### Google Search Agent

The Google Search Agent provides rapid, general-purpose information retrieval from the web. It is designed to access a wide array of online sources, including government and clinical websites, news articles, and other publicly available content.

To optimize performance, we design a parallel search and summarization architecture. This design addresses key challenges in web-scale data processing, namely: 1) maximizing search throughput; 2) preserving the finite context window of LLMs; and 3) mitigating performance degradation caused by irrelevant or excessively long text.

Our methodology is as follows: An Orchestrator generates multiple search queries, which are executed concurrently via the Google Search API. Each retrieved web page is then routed to a dedicated LLM instance. These instances operate in parallel to summarize their assigned content, guided by a prompt to focus on information relevant to the initial query. This strategy of parallelizing search and summarization significantly increases processing speed and serves as an effective filter, ensuring that only the most relevant information is retained. Finally, individual summaries are aggregated and synthesized into a consolidated report that includes robust source citations. Figure 16 illustrates the workflow of the Google Search Agent.

**Fig. 16.**
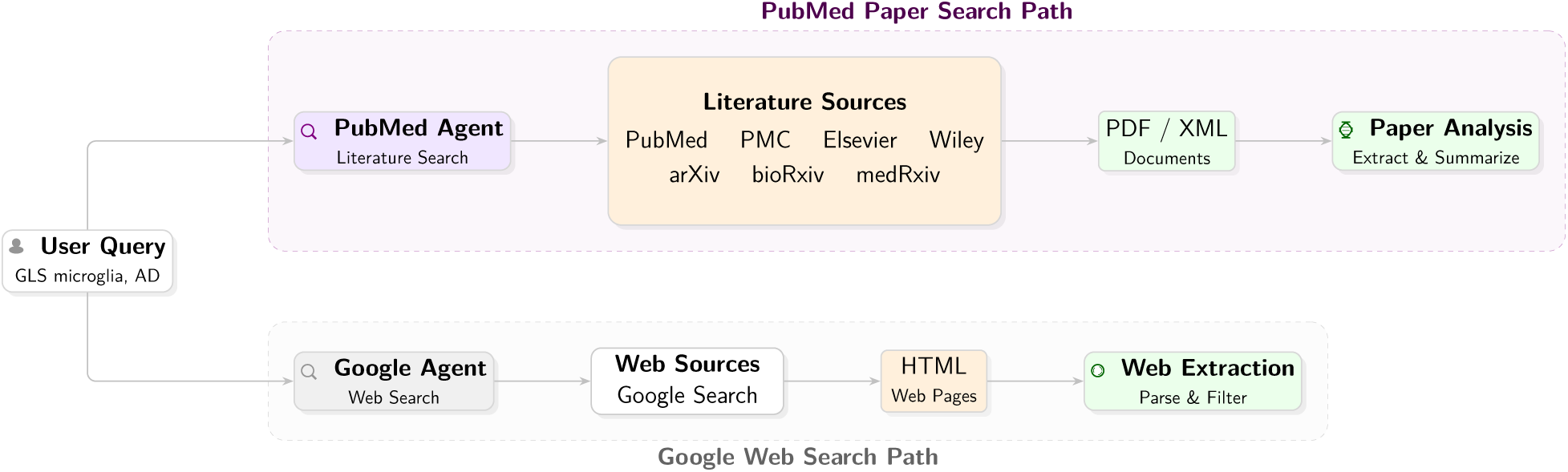
Overview of PubMed Paper Search Agent and Google Search Agent. **(Upper path)** The PubMed Paper Search Agent helps to find the latest and most domain-specific research and articles, delivering highly reliable, peer-reviewed scientific content ideal for evidence-based medical insights. **(Lower path)** The Google Search Agent helps to find the latest yet general information about the query, offering broad, up-to-date, and diverse information, including the latest developments not yet indexed in specialized databases.

### PubMed Paper Search Agent

The PubMedSearchAgent is a specialized agent designed to navigate and extract information from the biomedical scientific literature indexed within the PubMed database. Its primary function is to serve as a high-fidelity source for answering complex biomedical questions by retrieving peer-reviewed research articles, clinical studies, and systematic reviews.

The agent’s workflow begins when it receives targeted search queries from the Orchestrator. It utilizes the PubMed API to conduct a comprehensive search, returning a list of relevant publications with their DOIs. Using the list of DOIs, the agent attempts to download the full text of each article directly from the publisher’s platform. Our current implementation covers major scientific publishers and preprint servers, including PubMed Central (PMC), Elsevier, Wiley, arXiv, bioRxiv, and medRxiv, when their APIs are available. For each gene, the system can process information from up to 50 papers at reasonable speed.

We deploy a parallel extraction and summarization strategy to process the dense scientific content efficiently. Each article is assigned to a dedicated LLM instance. These LLMs are prompted to scrutinize the full text and identify and extract key evidence—such as methodologies, patient cohorts, statistical results, and author conclusions—that is directly relevant to the initial query. This parallel approach ensures that the agent can efficiently process large volumes of literature while isolating the most critical information.

The final output is a synthesized report that includes a summary of the findings from the retrieved articles, along with article titles. Figure 16 illustrates the workflow of the PubMed Paper Search Agent.

### BioMarkerKG Agent

The BioMarkerKG Agent is a specialized biomedical GraphRAG agent that retrieves biomedical subgraphs from a knowledge graph database. This fine-tuned graph language model processes biomedical queries and returns knowledge graph indices, generating subgraphs that contain enriched relationships between biomedical entities. This subgraph, which reveals disease mechanisms, drug-target interactions, or potential biomarkers, is then passed to a large language model (LLM) to synthesize a final, grounded response. Crucially, the BioMarkerKG Agent is invoked early in the workflow; its initial subgraph retrieval serves as a grounding framework that informs the search strategies of other agents, such as those for PubMed and web searches.

We construct the BioMarkerKG database using STaRK-Prime [58], a large-scale biomedical knowledge graph containing 129,375 entities and 8,100,498 relations across diseases, drugs, and genes/proteins. We convert entities and relations into textual embeddings alongside their descriptions and index the graph database for efficient subgraph retrieval.

We design a RAG QA fine-tuning task using 111,413 biomedical QA pairs from STaRK-Prime’s PrimeKG dataset. For example: Question: "I am looking for a gene or protein that plays a role in ribosomal operations and has an interaction with the protein RPL23. It should be linked to a disorder common with RPL23, and its absence should be a known cause of Diamond-Blackfan anemia. Which gene or protein fits these criteria?" Answer: "2496, 2977, 4933, 59, 717, 1518, 6384, 4789, 2137, 827, 1948, 349, 5054, 7423" (node entity indices).

We implement the training using a graph language model [18] specialized for RAG systems, comprising a graph encoder and a large language model. The graph encoder employs a Graph

Neural Network (GNN) architecture, specifically Graph Attention Networks (GAT) [52], paired with Llama-3.1-8B. This architecture effectively encodes graph structure and entity descriptions, enabling the model to understand retrieved subgraph structures and translate them into textual understanding. Fine-tuning on the RAG QA dataset teaches the model to map queries to corresponding entity indices in the graph database. Figure 18 illustrates the workflow of the BioMarkerKG Agent, and Figure 17 illustrates its information flow relative to an entire workflow realization.

**Fig. 17.**
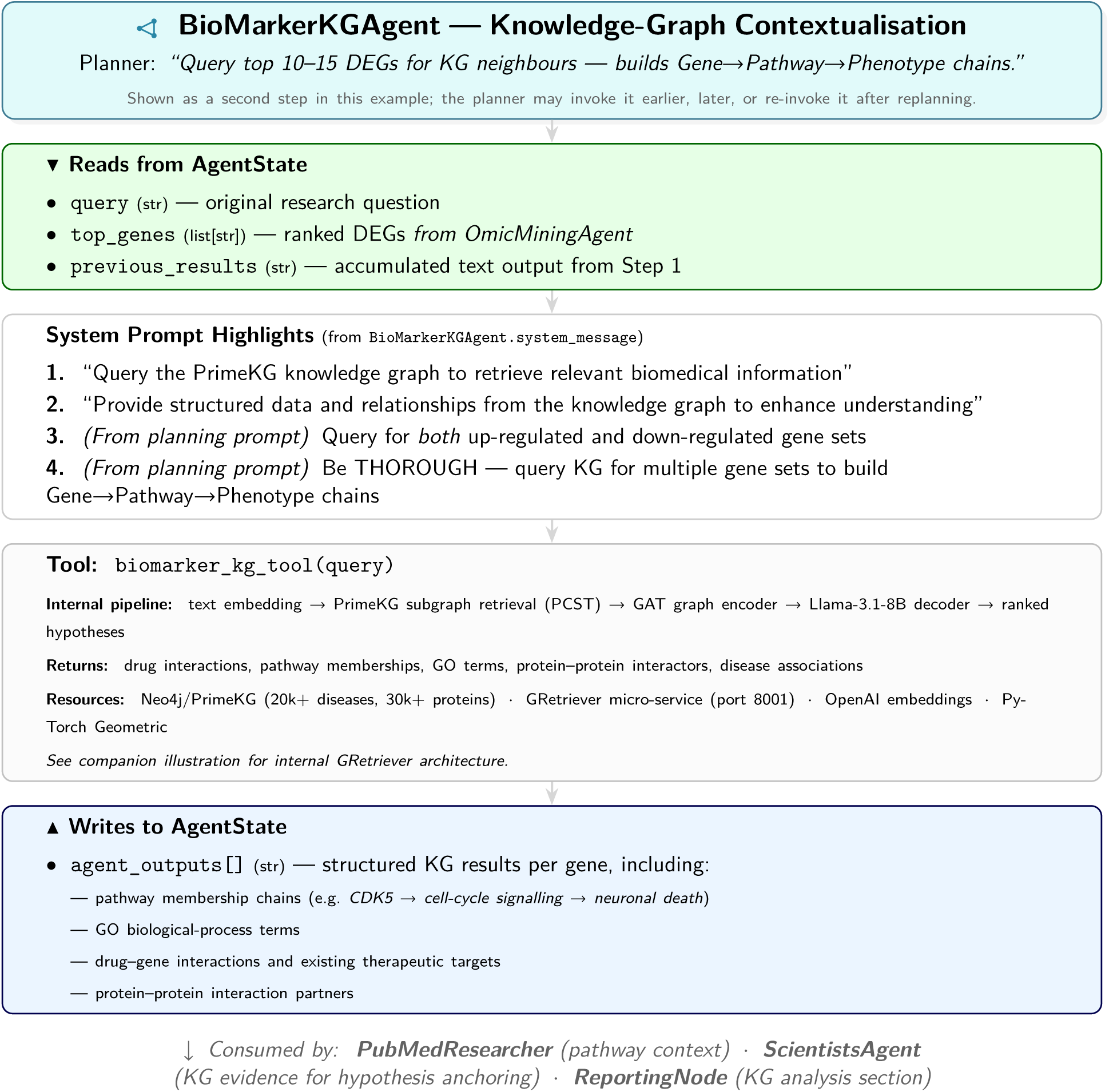
BioMarkerKGAgent. — workflow interface view. In the typical realisation shown here this agent runs second, but the LLM planner can reorder, skip, or re-invoke any agent at any point—for instance, the planner may call it again after literature search reveals new gene targets. When invoked, it reads the `top_genes` list from `AgentState` and queries the PrimeKG knowledge graph for each gene’s neighbourhood, targeting the top 10–15 DEGs to build explicit Gene→Pathway→Phenotype chains. Internally, the tool communicates with a GRetriever micro-service that extracts PCST subgraphs, encodes them with a Graph Attention Network, and decodes answers with a graph-adapted Llama-3.1-8B (see companion illustration for architecture details). Its output—appended to `agent_outputs[]`—provides the pathway and interaction context that downstream agents use to anchor hypotheses in structured biomedical knowledge.

**Fig. 18.**
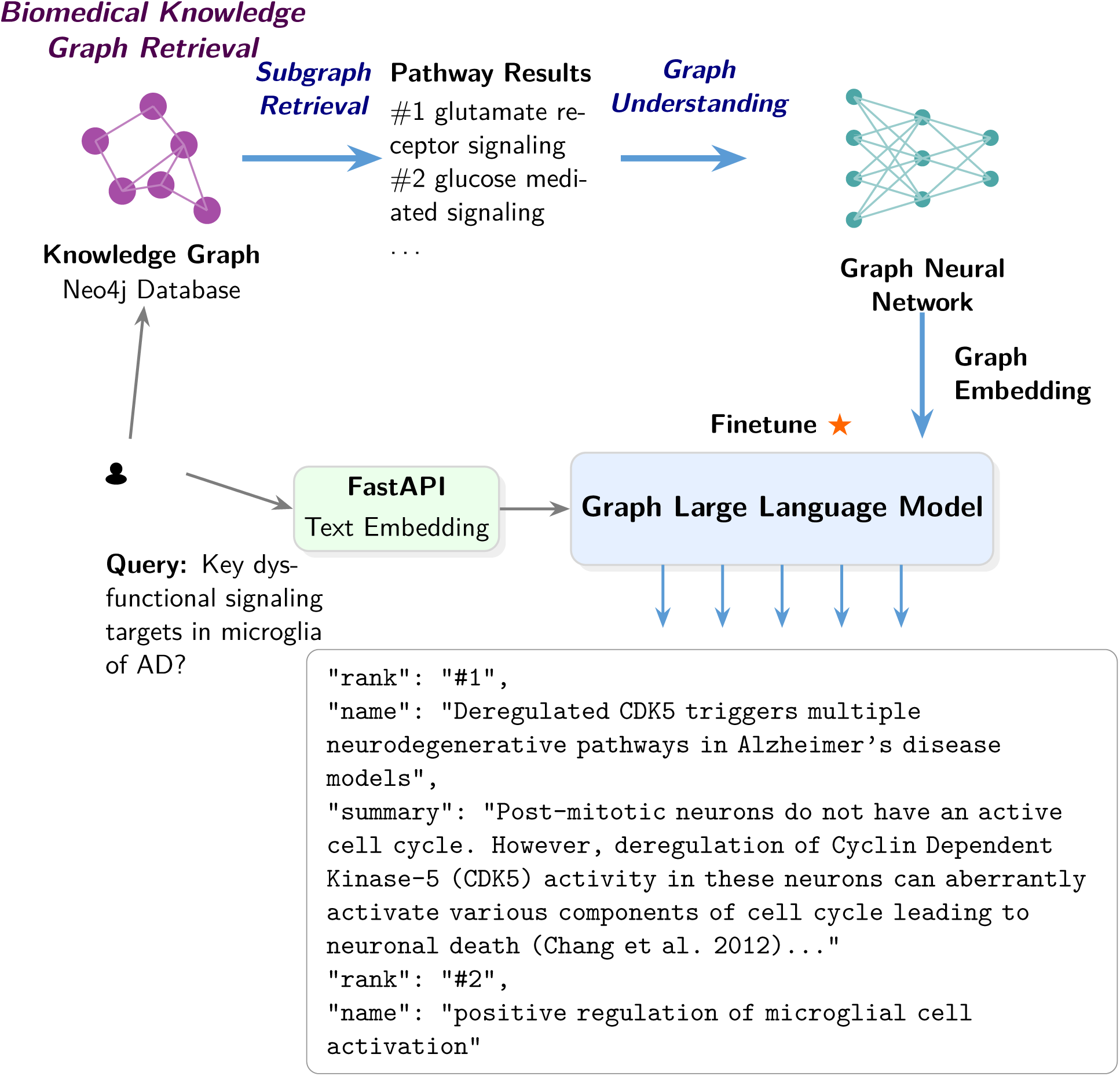
Overview of BioMarkerKG Agent. A retriever-specific language model—in this case a graph LLM—is trained to adapt to graph-based retrieval-based inferencing. When this agent gets a query, the text embedding of the query and the graph embedding of a knowledge graph subgraph retrieval result of the query are passed into the model simultaneously, providing both text and knowledge graph grounding for the results. The model is specifically fine-tuned to answer questions based on graph retrieval results.

### The Reporting Node and Appendix Building

Once every sub-task in the execution plan has been completed (or has exhausted its retry budget), the LangGraph state machine transitions to the Reporter Node—the terminal node of the orchestration graph. The Reporter Node is responsible for (1) synthesizing the accumulated evidence from all sub-agents into a single, coherent research report, (2) generating a structured appendix that preserves the full provenance of the analysis, and (3) compiling the output into both Markdown and PDF formats for downstream consumption.

Report synthesis. The Reporter Node collects all completed (and failed) tasks from the plan field of the `AgentState`, together with each task’s assigned agent and result summary stored in `agent_outputs`. These are assembled into a structured reporting prompt that prescribes a six-section output format: (1) Omics Data Analysis Summary—sample sizes, DEG counts, and a top-gene table ranked by FDR; (2) Knowledge Graph Analysis—first-neighbor entities categorized into drugs, pathways, GO terms, and diseases; (3) Literature-Validated Targets—the intersection of omics hits and literature evidence, with per-gene citations; (4) Pathway Enrichment Analysis—top KEGG, Reactome, and GO terms with embedded plots; (5) Gene-Anchored Mechanistic Hypotheses—three to five ranked hypotheses scored (MedHypoRank-GenePath) covering fit-to-evidence, mechanistic plausibility, testability, novelty, clinical impact, parsimony, and confounding risk, each accompanied by falsifiable predictions and a minimal validation experiment set with quantitative support/refute criteria; and (6) Conclusions with immediate next steps and full references. The system message accompanying the prompt enforces strict evidence standards: every claim must carry a DOI or PMID, and unsupported connections must be flagged explicitly rather than extrapolated. Because sub-agents save their visualization outputs—volcano plots, KEGG dotplots, enrichment bar charts—to predictable paths within the session directory (e.g. volcano_plots/, plots/, enrichment_results/), the Reporter is prompted to embed these figures using relative Markdown image syntax (e.g. ![Volcano Plot](volcano_plots/volcano_plot.png)). Since the compiled report resides in the same session directory as the generated plots, these relative references resolve correctly during PDF compilation, yielding a self-contained document with inline visualizations.

#### Appendix generation

After the LLM produces the main report text, the system programmatically appends a five-section appendix that captures the full execution provenance without relying on the LLM to recall or reconstruct it:

~~~
report += " \n\n " + build_execution_appendix (state) write_report (report, session_dir)
~~~

- **A1 — Session Information:** session identifier, generation timestamp, and the verbatim user query.
- **A2 — Execution Plan:** a table listing every sub-task with its task identifier, assigned agent, description (truncated to 60 characters for readability), and final status (completed, failed, or pending).
- **A3 — Process Log:** the last 50 entries of the timestamped `process_log`, recording planning decisions, per-task execution outcomes (success or failure with error summaries), replan triggers, and the reporting phase itself.
- **A4 — Shared Data Summary:** the structured data extracted programmatically during execution—up to 50 top DEGs with log2FC, FDR, and direction; disease and cell-type context; up to 20 literature DOIs; and up to 20 enriched pathway names.
- **A5 — Agent Outputs Summary:** the raw output from each sub-agent, keyed by task identifier and truncated at 3,000 characters per entry to balance completeness with readability.

This appendix serves a dual purpose. For the end user it documents exactly which datasets were queried, how many samples were analyzed, which literature sources were consulted, and what the intermediate findings were at each stage—providing the provenance chain necessary for scientific reproducibility. For system developers it supplies a diagnostic trace: if a hypothesis in the main report appears unsupported, the appendix reveals whether the upstream agent produced the expected gene list, whether the knowledge-graph query returned meaningful neighbours, or whether a PubMed search failed silently.

PDF compilation. The combined report-plus-appendix is first written as a Markdown file with YAML front matter (session ID, timestamp, query). The system then attempts PDF compilation via a three-strategy cascade: (1) `pandoc` with a L^A^T_E_X engine (`xelatex` or `pdflatex`), producing a table of contents, syntax-highlighted code blocks, and properly typeset tables; (2) if no L^A^T_E_X engine is available, `pandoc` renders to a styled HTML intermediate that `wkhtmltopdf` converts to PDF with custom CSS for image sizing and page-break control; (3) as a final fallback, `pandoc` produces a standalone HTML report styled via the Water.css stylesheet. All strategies set the session directory as the `resource-path`, ensuring that relative image references to plots and enrichment figures resolve correctly regardless of compilation method.

Conversation log. In addition to the report and appendix, the system persists a JSON conversation log containing the session identifier, the execution plan with per-task statuses, the full process log, truncated agent outputs (1,000 characters each), and the first 5,000 characters of the final report. This log enables continuation from a previous session’s state and facilitates cross-session comparison of agent behavior on different queries.

### Network Plot Generation and Validation

In addition to the Markdown/PDF report and the provenance appendix, the Reporter layer can emit a structured graph specification summarizing the main biological claims synthesized across the omics, knowledge-graph, literature, and translational-analysis branches. This artifact is not a raw knowledge-graph export and is not a direct omics visualization. Instead, it provides a reporter-derived integrative view of the final case narrative. In the current implementation, each network figure is associated with three files: a rendered PNG embedded in the report, a corresponding `*.spec.json` file containing the graph structure and styling instructions emitted by the Reporter, and a standalone `network_renderer.R` script that consumes the JSON specification and regenerates the final plot. This design separates LLM-based synthesis from deterministic rendering while preserving the intermediate representation.

The graph specification is emitted downstream of report synthesis and should be interpreted as a visual abstraction of the report’s core claims. The Reporter first integrates top omics signals, pathway-enrichment summaries, knowledge-graph neighbours, literature-supported mechanisms, and translational candidates into a ranked narrative, and then distills this narrative into graph nodes and edges representing anchor genes, contextual biology, and therapy-related hypotheses. Different edges therefore carry different semantics: some denote DEG-anchored functional associations, some denote contextual links supported by the knowledge graph or literature, and others denote biomarker-to-therapy vulnerability mappings. These figures should therefore be interpreted as integrative claim maps rather than as direct molecular interaction maps.

Because the network plot sits above the primary omics outputs, its validation proceeds at three levels. Structural validation checks that the Reporter-emitted JSON specification is complete, internally consistent, and renderable. Provenance validation checks that the main nodes and edges can be traced back to upstream evidence sources, including differential expression, pathway enrichment, knowledge-graph summaries, literature citations, and explicit statements in the revised report text. Biological review then distinguishes stable claim-supported edges from links that are merely plausible, literature-mediated, or visually stronger than the available evidence tier justifies. Within OCA, the network plot serves as a compact synthesis layer and as a practical interface for human review of the final multi-agent interpretation. For this reason, we treat it primarily as a Supplementary Information artifact unless it has undergone explicit manual curation and evidence-tier labeling.

### From Core Agent to User Application: A Deployment Architecture

The implementation of the agentic system does not deviate substantially from conventional approaches to deploying a hosted web-based chatbot. Indeed, the wealth of documentation, tutorials, and community-driven resources—both official and unofficial—renders the process relatively straightforward, provided that one adheres to established best practices. Modern software engineering benefits from a mature ecosystem of Python-based frameworks for the frontend, backend, database integration, and deployment pipelines, all of which can be leveraged to ensure robustness, modularity, and scalability. By closely following proven industry-standard solutions, we inherit the safety features and design principles that enable the system to be reliably delivered to the user.

In this project, the backend is constructed using the Flask [48] framework, which provides the essential routing mechanisms, event handling, and request/response management required to expose services to the user. Flask also facilitates seamless communication between the user interface and the underlying system state, enabling interactions with persistent entities such as the user and session tables. To manage these relational data structures, we employ SQLAlchemy [2], a powerful Object Relational Mapper (ORM) that abstracts raw SQL into high-level Python objects. This abstraction not only improves developer productivity but also enhances maintainability by enforcing consistency across database operations.

Additional backend utilities are supported through the Werkzeug [47] library, which underlies Flask itself and provides critical low-level components, including secure password hashing, request and response objects, and debugging tools. For handling production workloads, the system is deployed with `Gunicorn`, a robust WSGI HTTP server that enables horizontal scaling by distributing requests across multiple worker processes. This ensures responsiveness under increased load and forms the backbone of the production deployment architecture. To make the service externally accessible during development and lightweight testing phases, we utilize ngrok [44], which provides secure tunnels from a public URL to the locally hosted server. This allows the system to be bound to the endpoint, thereby making it accessible to external users without requiring a full production-grade containerized cloud infrastructure during early stages.

For creating interactive visualizations and dashboards, the system can optionally integrate Dash [36], which provides a reactive framework for building analytical web applications with minimal JavaScript overhead.

Although the technical scaffolding of the system aligns with established web-application practices, the more intricate challenge lies in orchestrating the lifecycle of agent sessions. Specifically, this involves defining events that trigger the initialization and termination of agents, establishing communication channels for sending and receiving messages, and implementing mechanisms for returning interactive visualizations generated by tool calls. These components are crucial for ensuring that the agentic system remains responsive, interpretable, and extensible, while maintaining a smooth user experience. The following sections describe in detail how these challenges are addressed and how the overall system is adapted for seamless agent interaction.

### System Architecture Layers

Figure 19 illustrates the hierarchical organization of the system architecture, decomposed into layers.

**Fig. 19.**
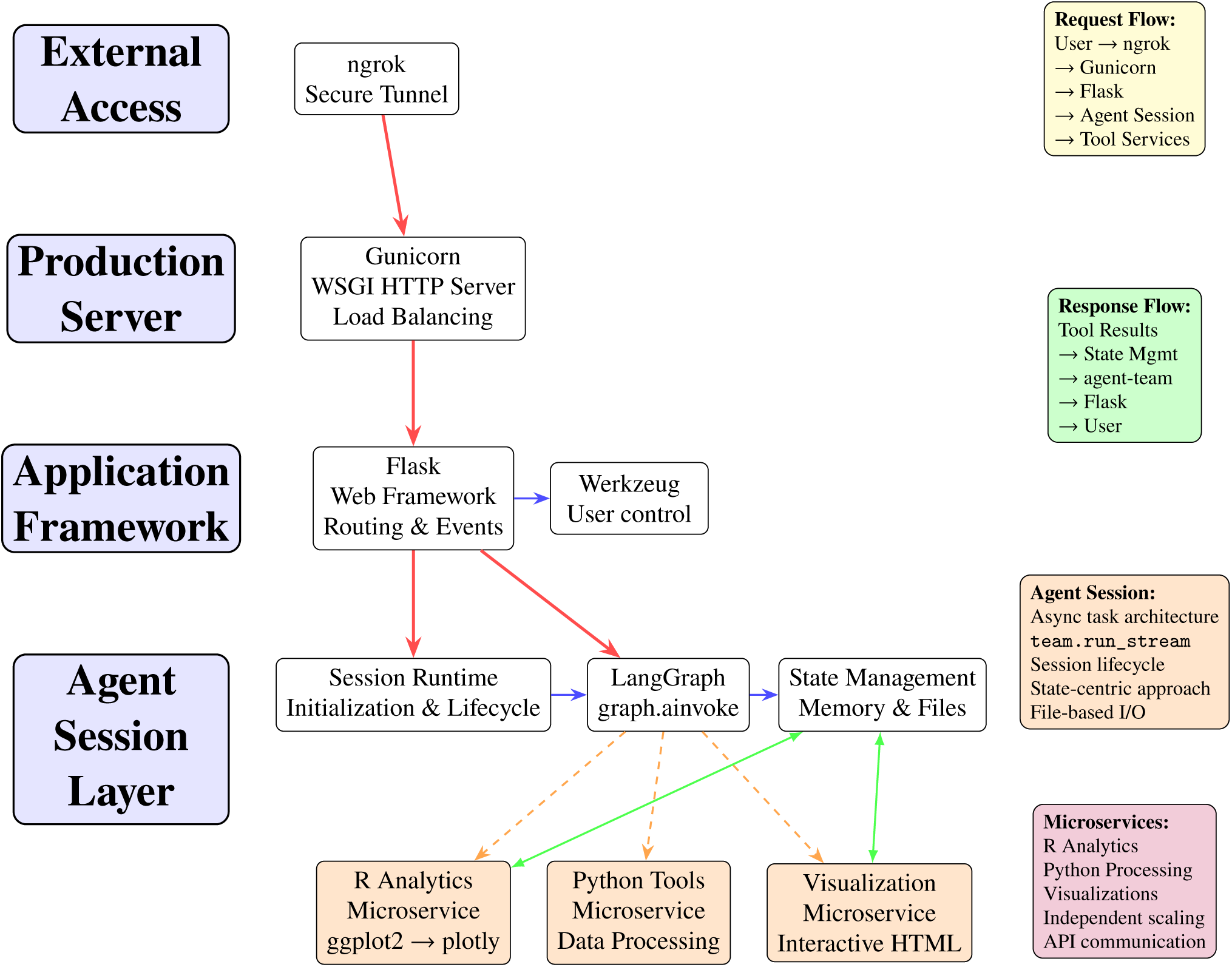
Service Architecture Diagram

### Agent Executor I/O

The agent executor orchestrates the entire agent execution lifecycle, typically embedded within sophisticated agentic frameworks that abstract away manual implementation details. These frameworks provide native communication channels and handle the intricate dynamics of agent interaction, eliminating the need for low-level process management. Each discrete event represents a state transition that furnishes the agent with contextual information and procedural guidance for subsequent actions. Consequently, maintaining a comprehensive state history effectively captures the agent’s cognitive trajectory—its reasoning patterns, decision-making processes, and execution steps.

**TABLE 1.**
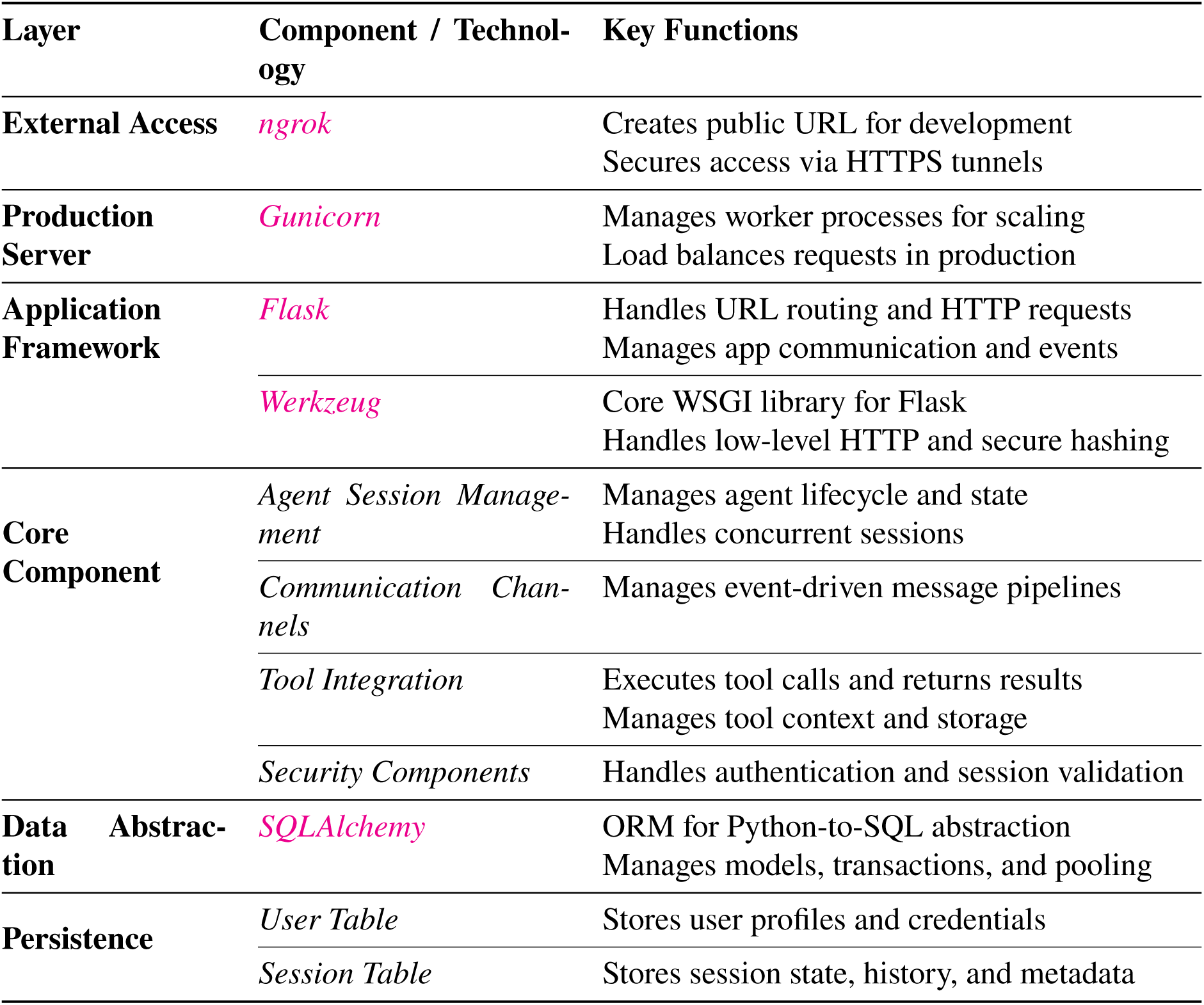
Concise System Architecture Layers and Components.

| Layer | Component / Technology | Key Functions |
| --- | --- | --- |
| External Access | *ngrok* | Creates public URL for development<br>Secures access via HTTPS tunnels |
| Production Server | *Gunicorn* | Manages worker processes for scaling<br>Load balances requests in production |
| Application Framework | *Flask* | Handles URL routing and HTTP requests<br>Manages app communication and events |
|  | *Werkzeug* | Core WSGI library for Flask<br>Handles low-level HTTP and secure hashing |
| Core Component | *Agent Session Management* | Manages agent lifecycle and state<br>Handles concurrent sessions |
|  | *Communication Channels* | Manages event-driven message pipelines |
|  | *Tool Integration* | Executes tool calls and returns results<br>Manages tool context and storage |
|  | *Security Components* | Handles authentication and session validation |
| Data Abstraction | *SQLAlchemy* | ORM for Python-to-SQL abstraction<br>Manages models, transactions, and pooling |
| Persistence | *User Table* | Stores user profiles and credentials |
|  | *Session Table* | Stores session state, history, and metadata |

This observation reveals an important architectural insight: despite the apparent complexity of multi-agent systems with their distributed components, hierarchical structures, intricate interaction protocols, and diagnostic mechanisms, the underlying operational logic of these systems can be distilled into state evolution processes. When we conceptualize agent interaction, reasoning, and problem-solving as functions operating on state transformations, we achieve a powerful abstraction that cuts through systemic complexity.

By anchoring our design philosophy in state-centric principles—where problem-solving emerges from state analysis and interactions are constructed around state manipulation functions—we capture the essential mechanics of agentic behavior. This paradigm shift from process-oriented to state-oriented thinking aims to yield remarkably clean, maintainable, and effective code architectures. The elegance comes from recognizing that beneath the surface complexity of distributed intelligence lies a unifying principle: agent behavior can be better understood and managed through disciplined state management.

In LangGraph’s implementation, this state-centric approach materializes through systematic control of the `graph.ainvoke` results, which encapsulate the dynamic execution state of the agent collective. File-based communication operates through directory synchronization mechanisms, enabling ‘FileReader’ components and text canvas memory systems to access shared resources efficiently. This architectural pattern crystallizes around two fundamental state domains: the evolving state of the agent team itself and the persistent state of the filesystem. These dual state repositories become the primary levers of control, transforming complex multi-agent orchestration into elegant state manipulation operations.

The outputs from `graph.ainvoke` are incremental—when new results become available, messages can be retrieved from this iterable. Direct printing provides a CLI interface, while parsing and triggering GUI updates yields a graphical interface. Results yielded in an iterable can also be serialized and saved to a folder incrementally to capture intermediate states. This enables failsafe features, resuming from a saved state, direct content inspection, or decoupling of agent and UI development. This approach is particularly beneficial when the agent has numerous dependencies or when execution is time-consuming or financially costly. The messages yielded from the team are native Python objects from LangGraph. Although sources are recorded and a parser can be made to show the results to the UI, we find it more efficient to maintain results in their native type. Rather than coercing results to strings and parsing the structure, directly handling Python types is significantly simpler and prevents faulty escape patterns that can lead to non-bijective encoding and decoding of hierarchical result structures.

There might be a hierarchical structure of objects returned, depending on the sources and types, so the parsing function should handle them case-by-case.

#### Visualizations: from ggplot to plotly to interactive UI

`ggplot2` [56] is a data visualization package for the R programming language. This framework enables the systematic creation of a wide array of statistical graphics by declaratively combining components such as data, aesthetic mappings, and geometric objects.

In our programmatic pipeline, static `ggplot` objects are converted into dynamic, interactive `plotly` objects. This conversion is accomplished using the ggplotly() function from the `plotly` R package [35].

The resulting `plotly` object can be exported as an HTML file. A key consideration is the method of including the necessary Plotly.js library.

- **Standalone HTML**: This method embeds the entire Plotly.js library directly within the HTML file. While this creates a self-contained document that functions offline, it results in a significantly larger file size.
- **CDN-linked HTML**: A more efficient alternative is to link to the Plotly.js library via a Content Delivery Network (CDN). This approach produces a lightweight HTML file that instructs the user’s browser to fetch the library from the CDN when the page is loaded. This method is ideal for embedding visuals into web-based UIs.

### Session Control

The agent runtime’s binding to Python’s awaitable task architecture reveals a deeper principle: computational agents are most naturally expressed as asynchronous processes rather than traditional thread-based entities. By encapsulating agent lifecycle management within a dedicated runtime creator class, we achieve precise control over initialization, execution, termination, and session enumeration. The superiority of asyncio-based session control over thread-centric approaches like psutil becomes evident when considering granularity—while psutil operates at the entire thread level and risks terminating the host application, async task management provides surgical precision over individual computational processes.

The execution flow embodies simplicity: a user message triggers agent team instantiation, which initiates the autonomous execution cycle. Termination occurs through either the agent executor’s intrinsic completion conditions or explicit task cancellation, both preserving the application integrity. This runtime creator class transcends mere process management by serving as the authoritative organizer of memory states and session identifiers. Through systematic recording and destruction of these ephemeral computational artifacts, the architecture promotes code cleanliness—each session becomes a self-contained, traceable, anddisposable computational unit.

Algorithm 1 presents the core session management logic.

### User Control

We build user control mechanisms to ensure that files belonging to one user do not interfere with the sessions and files of others. This creates a need for a user database, a backend to handle authentication events, and a way to connect that backend through to the agent executors. More importantly, we need to route session files to the correct location based on user ID and file storage system.

The authentication flow ensures proper isolation between user sessions while maintaining secure access control through the coordinated interaction of Flask [48], SQLAlchemy [2], and Werkzeug [47] components.

#### Algorithm 1 Agent Session Management with LangGraph Executor

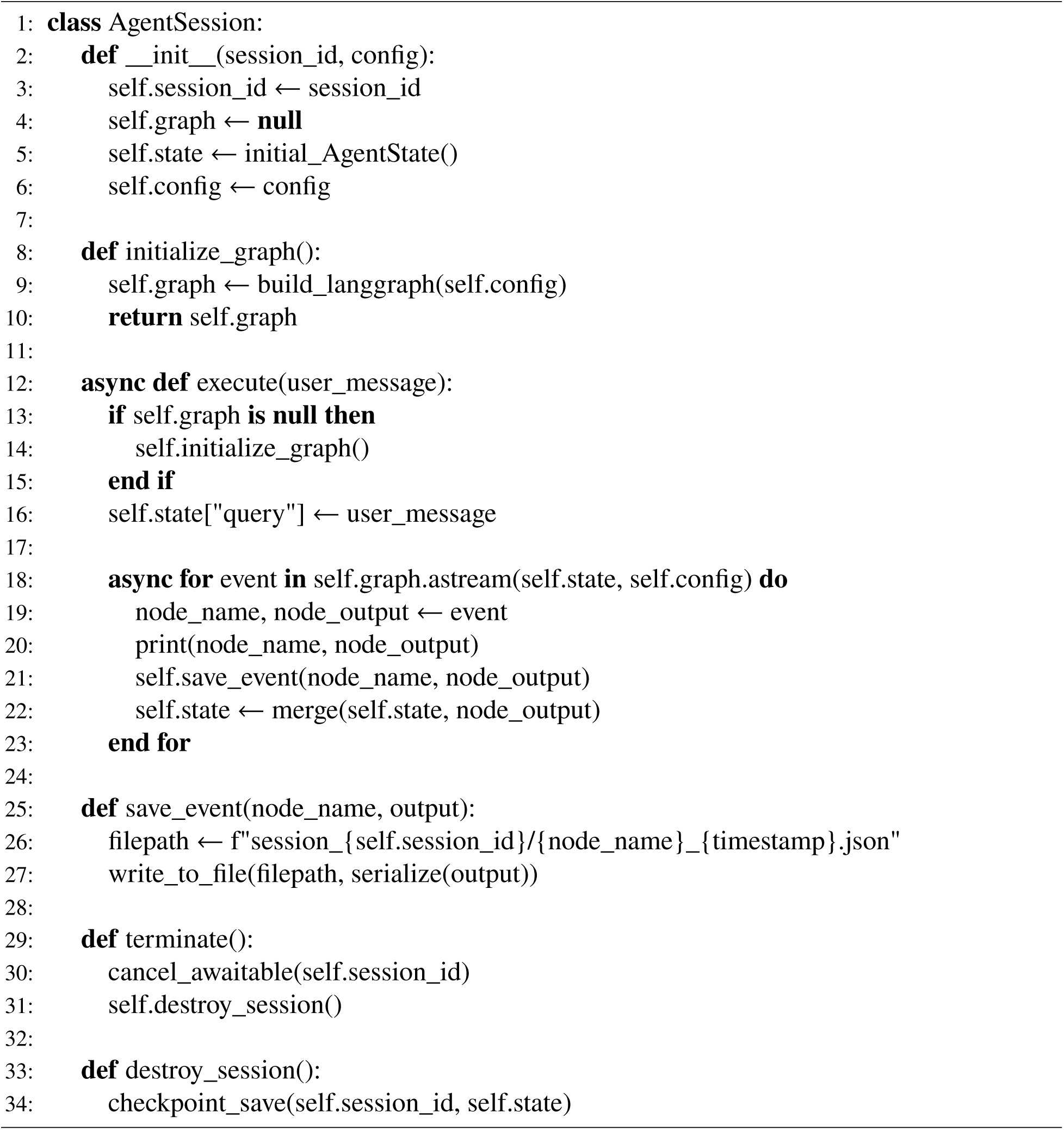

## ACKNOWLEDGMENTS

This research was partially supported by NIA 1R21AG078799-01A1, NIA 4R33AG078799, NINDS 1RM1NS132962-01, NLM 1R01LM013902-01A1, NIAID 1U19AI181984, and NIA R56AG065352.

